# Development of Enterovirus anti-viral agents that target the viral 2C protein

**DOI:** 10.1101/2022.10.06.511132

**Authors:** Rishabh Kejriwal, Tristan Evans, Joshua Calabrese, Lea Swistak, Lauren Alexandrescu, Michelle Cohen, Nahian Rahman, Niel Henriksen, Radha Charan Dash, M. Kyle Hadden, Nicola J. Stonehouse, David J. Rowlands, Natalie J. Kingston, Madeline Hartnoll, Samuel J. Dobson, Simon J. White

## Abstract

The enterovirus (EV) genus includes a number of important human and animal pathogens. EV-A71, EV-D68, poliovirus (PV), and coxsackievirus (CV) outbreaks have affected millions worldwide causing a range of upper respiratory, skin, neuromuscular diseases, including acute flaccid myelitis, and hand-foot-and-mouth disease. There are no FDA-approved anti-viral therapeutics for these enteroviruses. In this study, we describe novel broad spectrum anti-viral compounds targeting the conserved non-structural viral protein 2C that have low micro-molar to nanomolar IC_50_ values. The selection of resistant mutants resulted in amino acid substitutions in the viral capsid protein, implying a role for 2C in capsid assembly, as has been seen in PV. The assembly and encapsidation stages of the viral life cycle are not fully understood and the inhibitors reported here could be useful probes in understanding these processes.

## Introduction

Enteroviruses comprise a major group of viruses and include several important pathogens of humans and animals. They are small, non-enveloped, single-stranded positive-sense RNA viruses forming a genus within the *Picornaviridae* family^1^. Seven out of fifteen species of enteroviruses are human pathogens that cause various diseases of the skin, respiratory, circulatory, and nervous systems^2^. Enterovirus A71 (EV-A71) is one of the common causative agents of hand-foot-and-mouth disease. Although predominantly a disease in children, this can also affect adults and pose a serious threat to immunocompromised people. In a relatively small number of infections, EV-A71 can lead to more serious complications such as meningitis, encephalitis, and acute flaccid myelitis^3^ and in the Asia-Pacific region, EV-A71 outbreaks have had an overall mortality rate of 0.5 to 19 %^4^. The picornavirus EV-D68 is responsible for acute flaccid myelitis outbreaks in multiple countries, including the USA^5–8^, where there have been 682 confirmed cases since 2014^9^. Poliovirus (PV) causes paralytic poliomyelitis and although almost eradicated, it still poses a threat due to the reversion of attenuated vaccines to pathogenic vaccine-derived PV^10,11^, with cases recently reported in London and New York state. There are no FDA-approved drugs available against these viruses and there is an unmet clinical need for broad-spectrum anti-viral agents that show activity against enteroviruses.

The enterovirus 2C protein is an excellent target for the development of broad-spectrum antivirals. 2C is indispensable due to its multifunctional role in the viral life cycle including genome replication^12–15^, encapsidation of nascent RNA strands into new viral particles^16–20^, rearrangement of cellular membranes^21–24^, and facilitating the formation of capsid conformations required for efficient uncoating^25^. It is a highly conserved protein among enteroviruses (Figure S1 A) and shows very limited amino acid sequence identity to host proteins^26–28^. 2C has ATPase activity when oligomerized into a hexamer^29^ and can function *in vitro* as a AAA+ helicase^30^. Published crystal structures of an N-terminal truncated 2C lacking the first 115 amino acids of the 329-long protein from EV-A71, poliovirus, foot-and-mouth-disease virus, and hepatitis A virus show a conserved pocket^31–34^. This 2C pocket is adjacent to the ATP-binding site and is the site of interaction between the C-terminal alpha helix of one 2C subunit and another 2C subunit in the hexameric assembly. (Figure 1 A). The conservation of this pocket across EV 2C proteins makes this an attractive target for therapeutic development (Figure S1 B).

**Figure 1.**
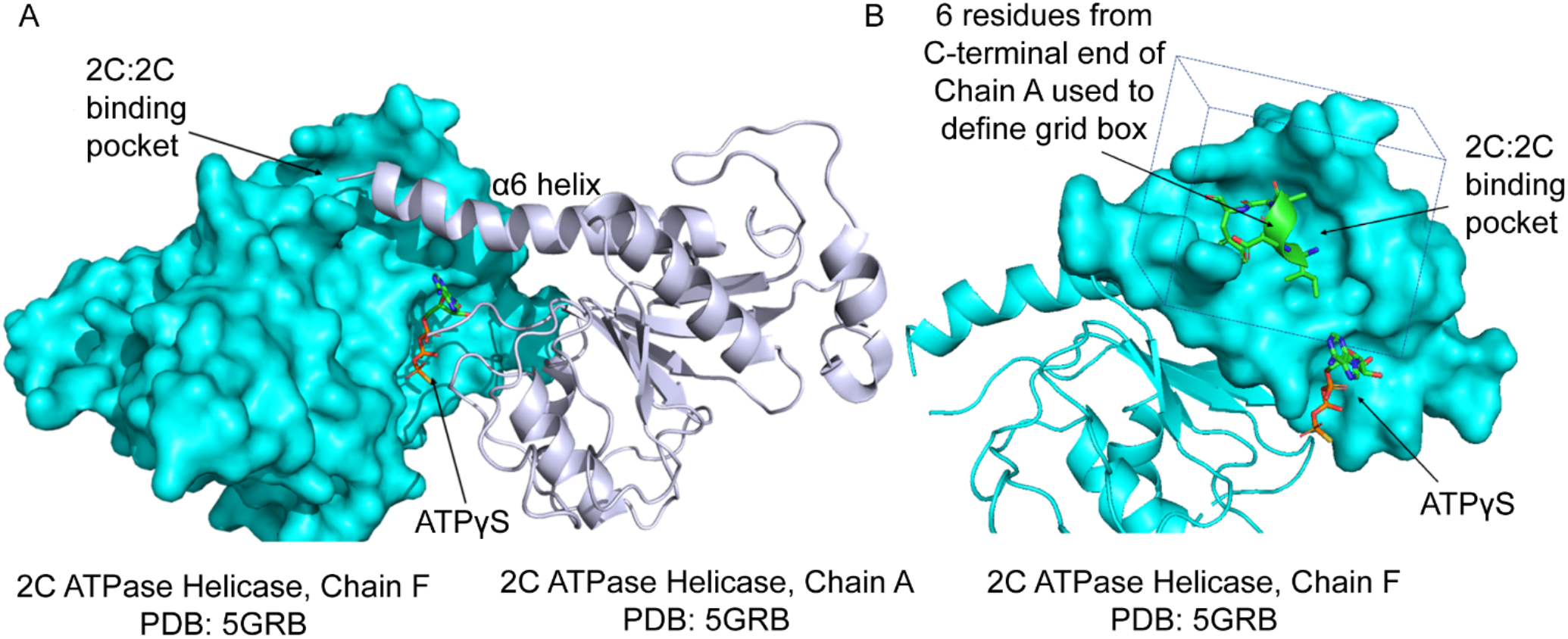
2C:2C binding pocket formed between 2C monomers. A) The surface rendition of Chain F (cyan) and the cartoon rendition of Chain A (white) from the crystal structure of EV-A71 2C (PDB:5GRB) are shown^31^. The α6 helix of Chain A is protruding into the adjacent 2C subunit (Chain F). The ATP molecule is shown as sticks. B) Part of the Chain F that forms the 2C:2C interacting pocket is shown as surface and the terminal six residues of Chain A C-terminal α6 helix are rendered as cartoon with sticks also shown. The box marks the pocket that was used for the *in silico* screen.

Here, we used a structure-based drug design approach to develop broad-spectrum anti-viral compounds that target the 2C pocket (Figure 1 B). We performed *in silico* screening of millions of commercially available small molecule compounds to identify those that were predicted to interact with the 2C pocket using the AtomNet® technology, a structure-based deep convolutional neural network virtual screening technology^35^. These small molecules were tested *in vitro* for 2C ATPase inhibition and in cells for anti-viral activity. Using these primary screening approaches and an extensive structure-activity relationship study, a potent compound, SJW-2C-227 with broad-spectrum anti-viral activity has been identified.

## Results

### *In silico* screening to identify potential 2C inhibitors

The partial crystal structure of EV-A71 2C was used for the structure-based drug discovery approach^31^. The crystal structure showed the C-terminal α6 helix of one monomer (PDB – 5GRB Chain A) protruding into a pocket in the adjacent 2C subunit (PDB – 5GRB Chain F) (Figure 1 A)^31^. Guan *et al*. suggested that this interaction was necessary for 2C oligomerization which is critical for its multi-functional role in the viral life cycle^31^. Further, on analyzing the 2C protein sequence from four Enteroviruses, it was seen that the residues forming the pocket were conserved (Figure S1 B), making this protein an attractive target for structure-based drug discovery. The terminal 7 amino acids (residues 323-329) of the C-terminal α6 helix from Chain A were used as the cognate ligand to select the residues forming the pocket (residues in Chain F). These residues in Chain F were L137, A138, G140, I141, R144, A145, D148, N277, F278, K279, R280, C281, S282, L284, and V285. These residues determined the screening grid box that was generated (Figure 1B). Using the AtomNet® technology, an *in silico* screening of a curated library of commercially available compounds was performed^35,36^. From this screen, 77 compounds were identified and tested in the primary assays for anti-viral activity.

### Two compounds show *in vitro* 2C ATPase inhibition and EV-A71 inhibition in cells

The 77 compounds identified from the *in silico* screen were tested in two primary assays, an ATPase assay to evaluate inhibition of 2C ATPase function and a cell-based assay to study the anti-viral activity of the compounds.

The ATPase assay is a colorimetric assay that uses malachite green to quantify the released inorganic phosphate following ATP hydrolysis by 2C^30^ (Figure S2). Successful inhibition of 2C reduces the production of inorganic phosphate from ATP hydrolysis, resulting in a decrease in absorbance. The 77 compounds were tested in this ATPase assay at a single concentration of 50 µM using the truncated Δ2C^116-329^. The truncated 2C protein was used for assays since it has been shown to be far more soluble than the full-length 2C protein^31,32^. We elected to use a cutoff of 3x standard deviation (3SD) below the average absorbance readout to identify 2C ATPase inhibitors. Four compounds, SJW-2C-1, SJW-2C-14, SJW-2C-33, and SJW-2C-69, inhibited the ATPase function of the Δ2C^116-329^, as indicated by the decrease in absorbance below the 3SD cutoff value of 2.53 for the DMSO-treated control (Figure 2 A). These four compounds were then studied further by performing a dose-response analysis. The IC_50_ value for this ATPase inhibition was calculated to be in the range of 7 µM to 70 µM (Figure 2 B). Previously, guanidinium hydrochloride (GnHCl) and fluoxetine have been shown to inhibit the ATPase activity of 2C^37,38^ and were used as positive controls here. The IC_50_ value for GnHCl was 83.58 mM while that of the racemic mixture of fluoxetine was 537.4 µM (Figure 2 C). These values are similar to previously published results, validating the assay conditions and the robustness of the assay. ^37–39^

**Figure 2.**
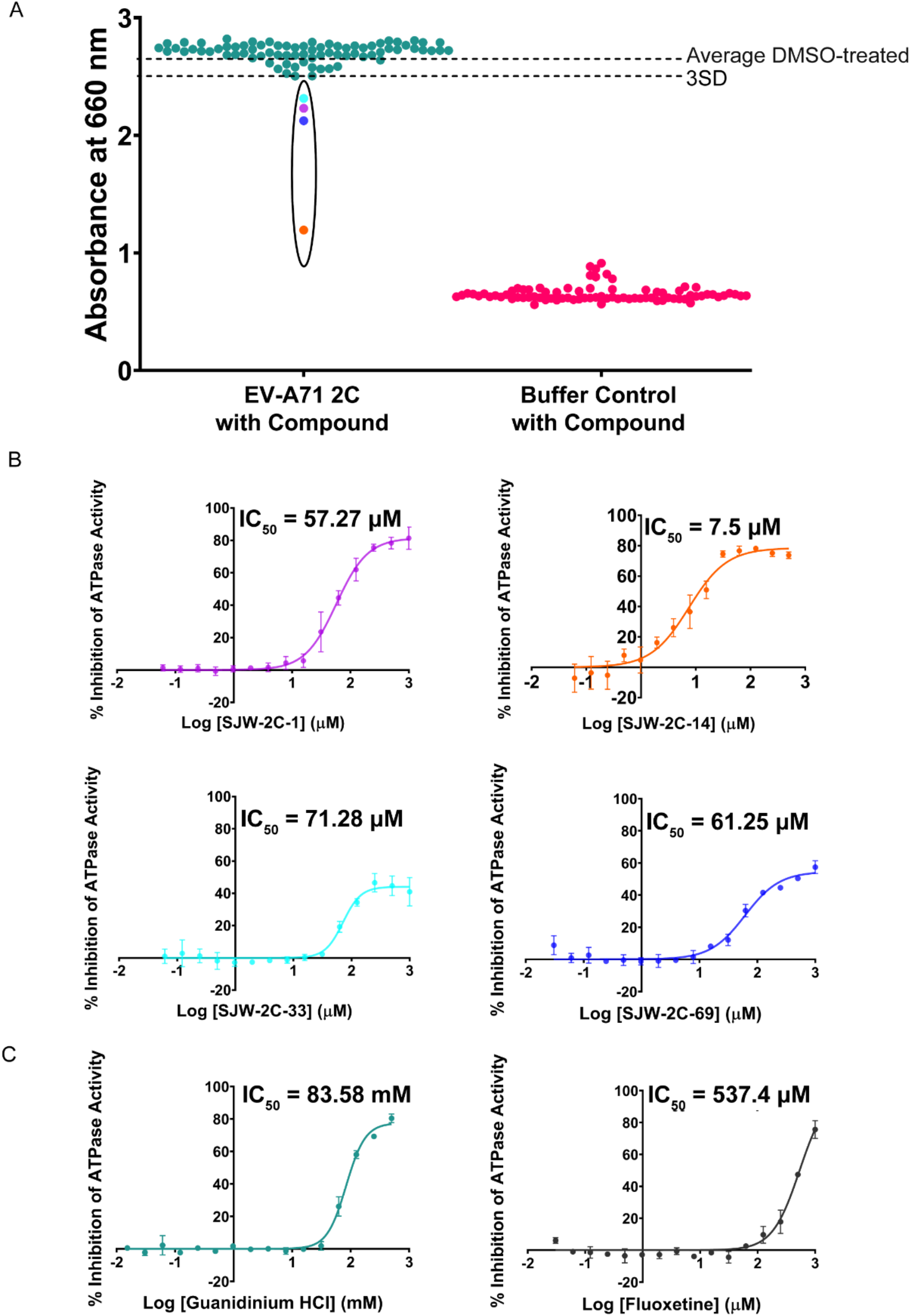
Compounds tested using the *in vitro* 2C ATPase inhibition assay. A) 77 compounds were screened to evaluate whether they were able to inhibit the ATPase function of Δ2C^116-329^ using the colorimetric malachite green-based assay at a final concentration of 50 µM. A decrease in absorbance at 660 nm indicates inhibition of the ATPase function. Four compounds inhibited Δ2C^116-329^ ATPase > 3SD below DMSO-treated control. B) Dose-response curves for SJW-2C-1, SJW-2C-14, SJW-2C-33, and SJW-2C-69 were performed to determine the IC_50_ between 0.06-1000 µM. C) Dose-response curves to determine the IC_50_ values for the two positive controls, guanidinium HCl (GnHCl) and fluoxetine, were performed. The concentration range for GnHCl was 0.015 mM to 500 mM and 0.06 µM to 1000 µM for fluoxetine. Graphed mean and SD, n=3.

A cell-based assay was performed to assess the anti-viral activity of the 77 compounds against EV-A71 in Vero cells. In this assay, luminescence decreases as a consequence of the cell destruction by virus induced cytopathic effects (CPE) and so increased luminescence indicates inhibition of viral replication. The 77 compounds were tested at a single concentration of 50 µM for anti-viral activity. A luminescence reading of 3SD above the average luminescence produced by infected cells (Cells+DMSO+Virus) was used as a threshold to identify anti-viral compounds. The cytotoxicity of the compounds was also tested and was reflected in the Cells+Drug wells. Two independent repeats were undertaken (Figure S3). Compounds that resulted in a luminescence reading above the 3SD cutoff in the Cells+Drug wells and the Cells+Drug+Virus wells were selected and using this criterion, 21 of the 77 compounds showed anti-viral activity (Figure 3 A). Out of the four ATPase inhibiting compounds, SJW-2C-14 (orange circle) and SJW-2C-69 (blue circle), inhibited EV-A71 while showing limited toxicity. SJW-2C-14 had a CC_50_ of 422.4 µM and SJW-2C-69 had a CC_50_ of > 400 µM. Using the IC_50_ and the CC_50_ values, we determined the selectivity index for SJW-2C-14 and SJW-2C-69 to be 9.44 and > 16.8 respectively (Table 1). The other two compounds identified in the *in vitro* ATPase assay, SJW-2C-1 (purple circle) and SJW-2C-33 (cyan circle), were cytotoxic as seen in the Cells+Drug wells and so were not considered for further testing and development (Figure 3 A). SJW-2C-14 and SJW-2C-69 were then evaluated using a dose-response study to calculate the IC_50_ values. Treatment with SJW-2C-14 or SJW-2C-69 inhibited the virus-induced CPE with a clear dose-dependent response (Figure 3 B). For this assay, we used GnHCl and dibucaine as positive controls since fluoxetine was ineffective against EV-A71 at non-toxic concentrations^40^. GnHCl and dibucaine are active against EV-A71 in cells with low mM and low µM IC_50_ values respectively^41^. In our assays, GnHCl inhibited EV-A71 with an IC_50_ of 0.14 mM while dibucaine treatment resulted in an IC_50_ of 4.5 µM. These values are similar to previously published values^41–45^, validating the experimental conditions and the robustness of the assay (Figure 3 C).

**Figure 3.**
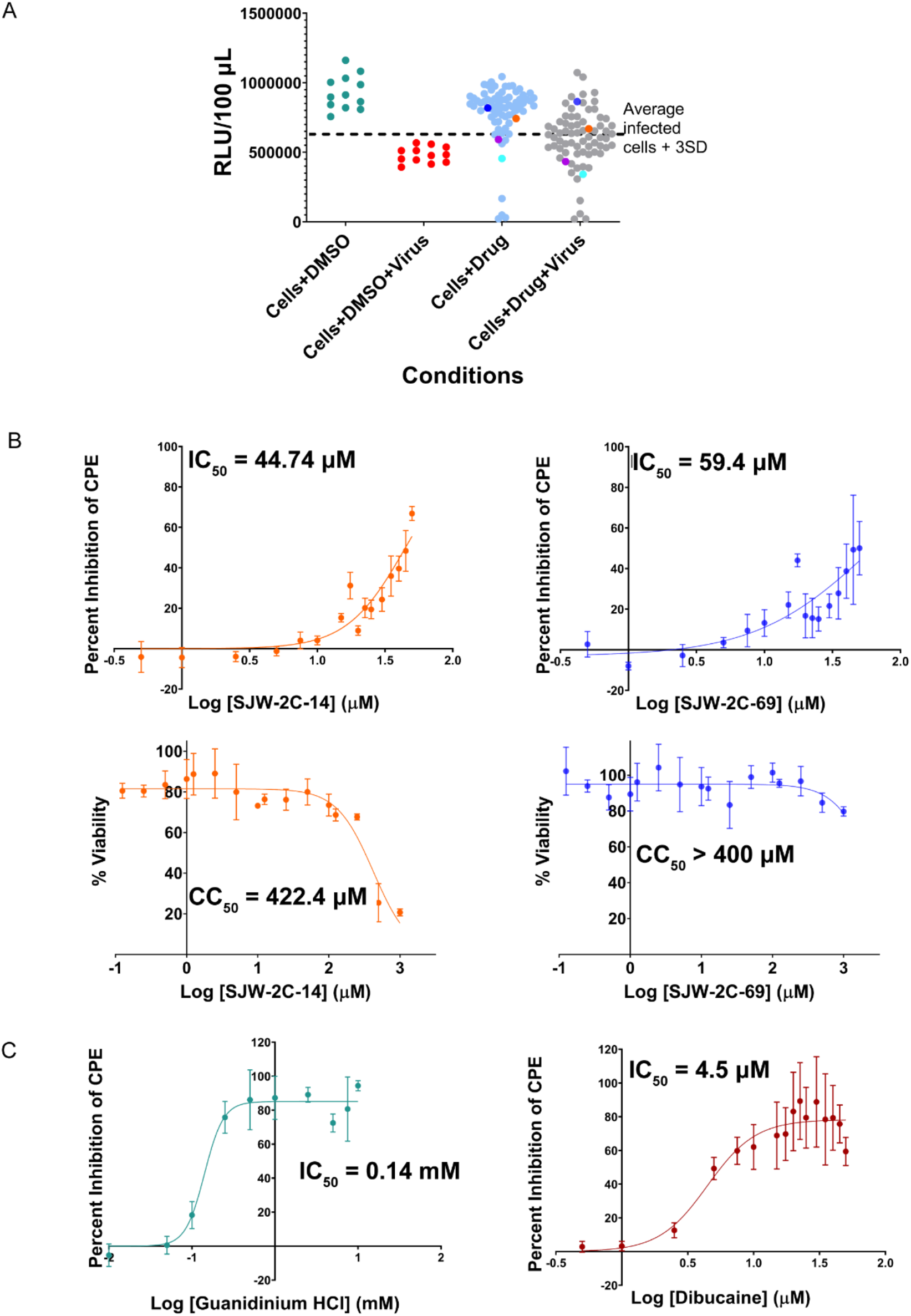
Compounds tested for inhibition of cytopathic effects (CPE) caused by EV-A71 in Vero cells and cytotoxicity. A) Compounds were screened at a single drug concentration of 50 µM. The Cell Titer Glo kit from Promega was used to read out the cell viability which directly correlates to the inhibition of CPE caused by EV-A71. A threshold of 3x standard deviation (3SD) above the average relative luminescence units (RLU) readout of the infected cells (Cells+DMSO+Virus) was used to identify the anti-viral compounds. The reaction volume was 100 µL. SJW-2C-1 is shown as purple circle, SJW-2C-14 is shown as orange circle, SJW-2C-33 is shown as cyan circle, and SJW-2C-69 is shown as blue circle. B) IC_50_ values for the anti-viral activity of SJW-2C-14 and SJW-2C-69 were calculated using dose-response curves and the concentration range was 0.5 µM to 50 µM. C) Cytotoxicity in Vero cells was determined for these two compounds using dose-response curves. The dosage range was from 0.125 µM to 1000 µM. D) Dose-response curves for the two positive controls, guanidinium HCl and dibucaine, were generated to determine their IC_50_ values. The concentration range for GnHCl was 0.01 mM to 10 mM and for dibucaine it was 0.5 µM to 50 µM. Colors as used in Figure 2. Graphed mean and SD, n=3.

**Table 1.** Antiviral activity of active compounds against EV-A71. The assay was performed three times from which the 95 % confidence interval range for IC_50_ was calculated using profile likelihood asymmetrical confidence intervals.

| **Compound Name** | **IC<sub>50</sub> (μM)** | **95 % confidence interval range (μM)** | **CC<sub>50</sub> (μM)** | **Selectivity Index** |
| --- | --- | --- | --- | --- |
| SJW-2C-14 | 44.74 | 41.05 to 50.16 | 422.4 | 9.44 |
| SJW-2C-69 | 59.4 | 44.25 to 96.63 | > 1000 | > 16.8 |

### Hit expansion to optimize SJW-2C-14 and SJW-2C-69

Having identified two compounds from the initial *in silico* screen that showed inhibition of 2C ATPase function and inhibition of the virus in a cell-based assay with limited toxicity, we next performed a round of hit expansion to identify structure-activity relationships (SAR) for our two compounds (SJW-2C-14 and SJW-2C-69) to improve the activity of the compounds (lower IC_50_ values) as well as their cytotoxicity (higher CC_50_ values). We focused on analog compounds that were commercially available, so-called “analog-by-catalog”.

Comparing the structures of SJW-2C-14 and SJW-2C-69 showed a central pharmacophore of a bicyclic ring (Figure 4 A, blue circle), similar to the benzimidazole of ATP. However, previous studies testing benzimidazole-containing compounds against enteroviruses did not show inhibition of the ATPase function of 2C^42,46^. To minimize ATP mimicking compounds, we avoided analogs with R-groups at positions equivalent to the N^6^ and the N^9^ position of the ATP molecule (orange circle and red circle respectively) (Figure 4 A). Instead, R-groups at the N^8^ position (green circle) and the 2-position were selected in the analogs to optimize binding to the 2C pocket. To explore this pharmacophore (Figure 4 B), we performed a geometric-based similarity study using binary fingerprints and selected 45 analogs from an in-house library, and tested them in the cell-based CPE inhibition assay against EV-A71 using Vero cells. Only those compounds that showed luminescence above the 3SD value were considered active. Of the 45 analogs, six compounds were active (Figure 4 C and S4). Upon determining their IC_50_ and CC_50_ values using dose-response curves, SJW-2C-184 showed the best activity, with an IC_50_ of 2.9 µM (Figure 4 D). It showed no toxicity to Vero cells up to 200 µM resulting in a selectivity index of > 68.9 (Figure 4 E and Table S1).

**Figure 4.**
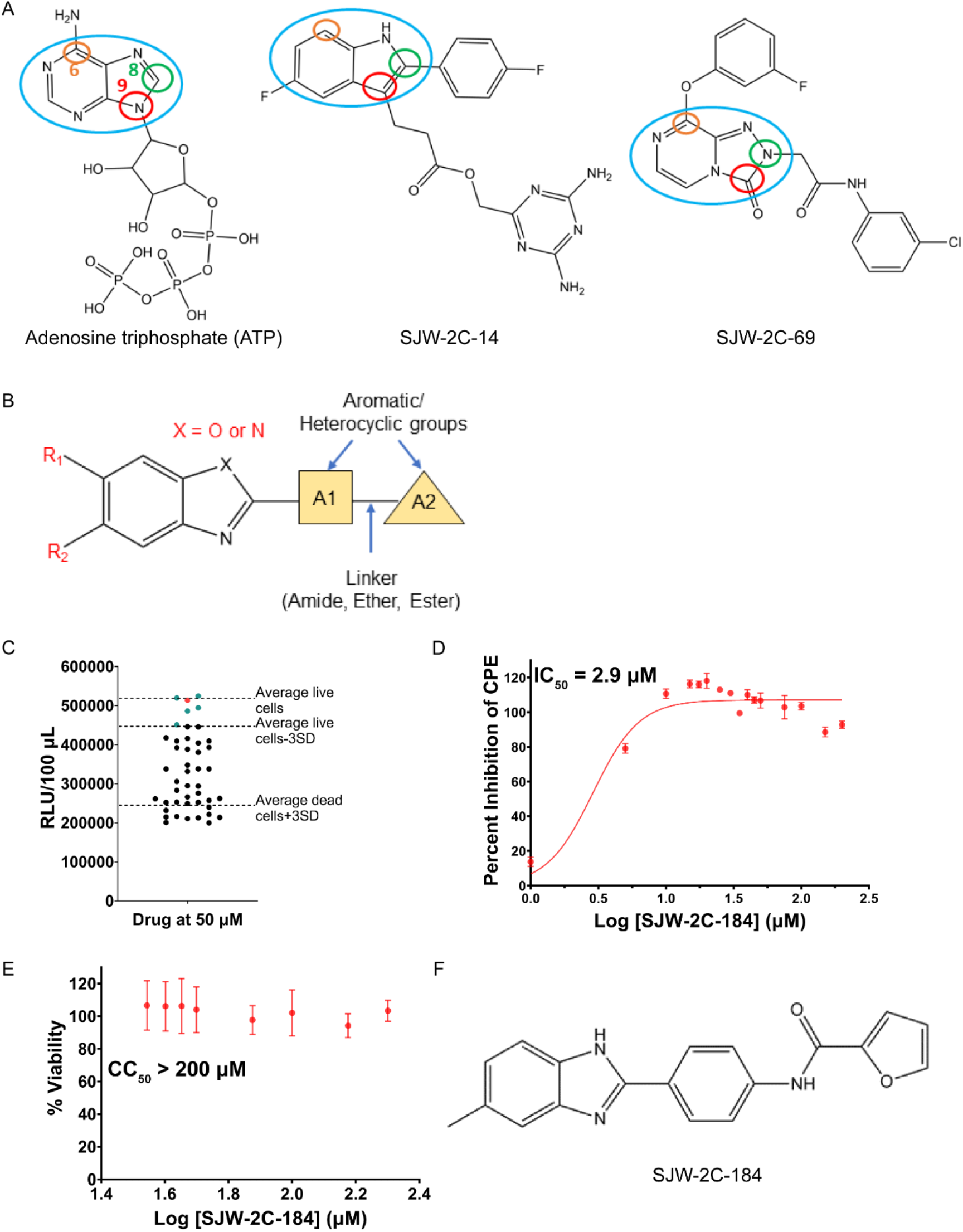
Structure-activity relationship studies. A) Comparison of the chemical structures of adenosine triphosphate (ATP), SJW-2C-14, and SJW-2C-69 showing that by including R-groups at the N^8^ position (green circle) and by avoiding R-groups at the N^6^ (orange circle) and N^9^ (red circle) positions, we could select analogs that were different from ATP. B) Pharmacophore model developed to select analogs from the in-house library. The central ring is either a benzimidazole (if X = N) or benzoxazole (X = O) with either R_1_ or R_2_ or both R groups. Further, the molecule can either have A_1_ or both A_1_ and A_2_ aromatic/heterocyclic linked by an amide or ether or ester linker. C) 45 analogs of SJW-2C-14 and SJW-2C-69 were tested for CPE inhibition in the cell-based assay at a final concentration of 50 µM. Six compounds showed relative luminescence units (RLU) reading above the 3SD cutoff and are shown as either green circles or a red circle and the ones that were below the cutoff are shown as black circles. SJW-2C-184, the analog with the best activity, is shown as a red circle for better visualization. The reaction volume was 100 µL. D) Dose-response curve for SJW-2C-184 was carried out to determine its IC_50_. The dose range for this assay was 1 µM to 200 µM. Graphed mean and SD, n=3. E) CC_50_ of SJW-2C-184 was determined in Vero cells. The dose range for this assay was 35 µM to 200 µM. Graphed mean and SD, n=3. F) The chemical structure of SJW-2C-184.

Next, we conducted another round of hit expansion to identify further structure-activity relationships (SAR) for SJW-2C-184 (Figure 4 F). 67 analogs of SJW-2C-184 were identified to fit the pharmacophore structure and these were grouped into 4 categories, based on the pharmacophore model (Figure 4 B and Tables S2-S6). The 67 compounds were first tested for anti-viral activity in Vero cells against EV-A71 using the CPE inhibition assay. 40 analogs had a luminescence reading above the 3SD cutoff and were deemed active against the virus (Figure 5 A). We generated dose-response curves to determine the respective anti-viral IC_50_ values and CC_50_ values to analyze cytotoxicity. The antiviral IC_50_ values ranged from 0.7 µM to 250 µM with 9 analogs having IC_50_ ≤ 5 µM (Table 2, and S2-S6). The 67 analogs were also tested for Δ2C^116-329^ ATPase inhibition capability using the malachite green-based assay. 14 analogs inhibited the ATPase function of Δ2C^116-329^ (Figure 5 B). Eleven analogs showed anti-viral activity in cells as well as inhibition of the ATPase function of Δ2C^116-329^ (Table 3). SJW-2C-227 inhibited the virus-induced CPE with an IC_50_ of 2.66 µM (Table 2) and inhibited the ATPase function of Δ2C^116-329^ with a clear dose-dependent response (Table 3). Further, it had a CC_50_ value of 78.71 µM giving it a selectivity index of 29.6 (Figure 5 C).

**Figure 5.**
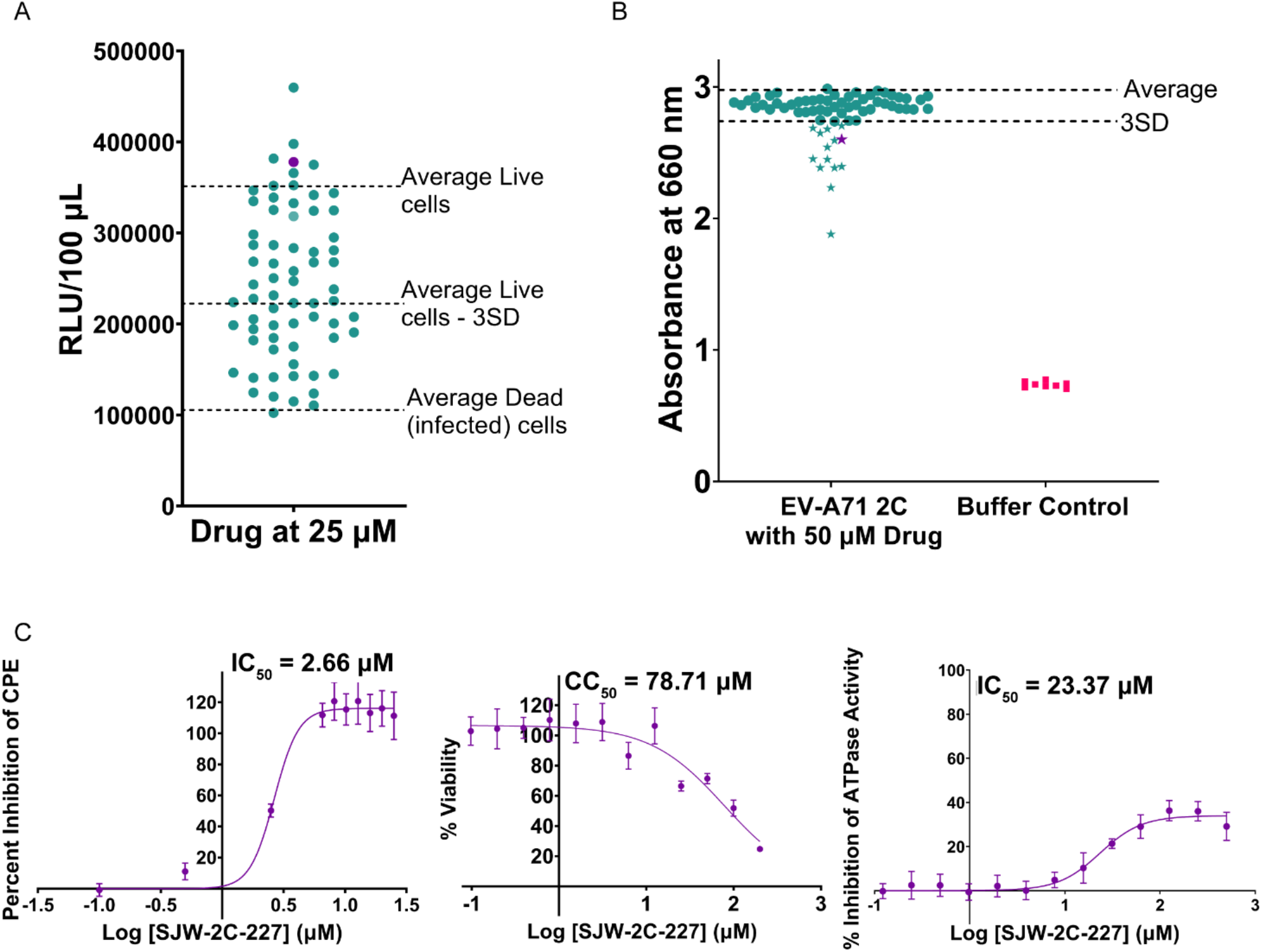
SAR screening of the 67 analogs of SJW-2C-184 using the cell-based assay and the *in vitro* ATPase assay. A) The 67 analogs were tested in the cell-based CPE inhibition assay at 25 µM. Promega Cell Titer Glo kit was used for this assay and higher relative luminescence units (RLU) indicates higher inhibition of the CPE caused by the virus. The reaction volume was 100 µL. B) Δ2C^116-329^ ATPase inhibition assay was carried out to evaluate whether these compounds can inhibit the ATPase function of Δ2C^116-329^ using the colorimetric malachite green-based assay at a final compound concentration of 50 µM. The stars highlight the compounds that showed activity in both the assays. C) Dose-response curves for SJW-2C-227 were generated to calculate IC_50_ value for antiviral activity (CPE inhibition) and Δ2C^116-329^ ATPase inhibition activity, and the CC_50_ value for the cytotoxicity. Graphed mean and SD, n=3.

**Table 2.**
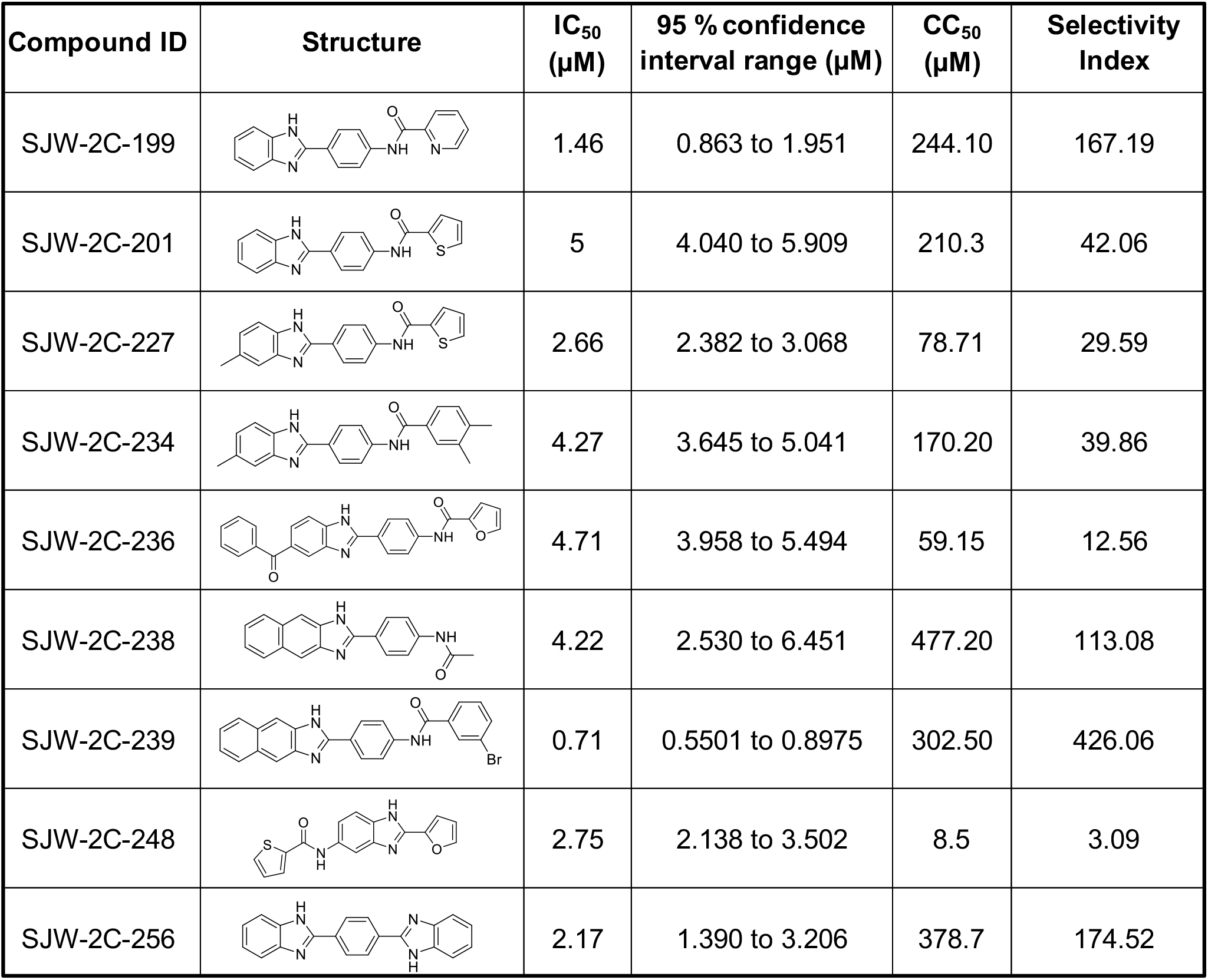
Analogs inhibiting virus-induced cellular cytopathic effect. The chemical structures of the top nine compounds with IC_50_ values ≤ 5 µM are shown below. The assay was performed three times from which the 95 % confidence interval range for IC_50_ was calculated using profile likelihood asymmetrical confidence intervals. Selectivity Index = CC_50_/IC_50_.

| Compound ID | Structure | IC<sub>50</sub> (μM) | 95 % confidence interval range (μM) | CC<sub>50</sub> (μM) | Selectivity Index |
| --- | --- | --- | --- | --- | --- |
| SJW-2C-199 |  | 1.46 | 0.863 to 1.951 | 244.10 | 167.19 |
| SJW-2C-201 |  | 5 | 4.040 to 5.909 | 210.3 | 42.06 |
| SJW-2C-227 |  | 2.66 | 2.382 to 3.068 | 78.71 | 29.59 |
| SJW-2C-234 |  | 4.27 | 3.645 to 5.041 | 170.20 | 39.86 |
| SJW-2C-236 |  | 4.71 | 3.958 to 5.494 | 59.15 | 12.56 |
| SJW-2C-238 |  | 4.22 | 2.530 to 6.451 | 477.20 | 113.08 |
| SJW-2C-239 |  | 0.71 | 0.5501 to 0.8975 | 302.50 | 426.06 |
| SJW-2C-248 |  | 2.75 | 2.138 to 3.502 | 8.5 | 3.09 |
| SJW-2C-256 |  | 2.17 | 1.390 to 3.206 | 378.7 | 174.52 |

**Table 3.** Analogs inhibiting 2C ATPase activity. IC_50_ values for ATPase inhibition were determined for analogs that inhibited EV-A71 in Vero cells and are as stated below. SJW-2C-194, SJW-2C-214, and SJW-2C-232 inhibited 2C ATPase but were not active in cells. The assay was performed three times from which the 95 % confidence interval range for IC_50_ was calculated using profile likelihood asymmetrical confidence intervals.

| **Scaffold #** | **Compound ID** | **IC<sub>50</sub> (μM)** | **95 % confidence interval range (μM)** |
| --- | --- | --- | --- |
| Scaffold 1A | SJW-2C-203 | 652.1 | 452.6 to 1076 |
|  | SJW-2C-211 | 18.06 | 10.37 to 47.73 |
|  | SJW-2C-219 | 0.55 | 0.3392 to 1.239 |
| Scaffold 1B | SJW-2C-227 | 23.37 | 17.67 to 30.99 |
|  | SJW-2C-228 | 20.59 | 15.56 to 29.18 |
|  | SJW-2C-231 | 37.26 | 31.21 to 46.08 |
|  | SJW-2C-234 | 55.04 | 40.31 to 85.50 |
|  | SJW-2C-235 | 18.96 | 15.26 to 23.91 |
|  | SJW-2C-237 | 17.35 | 12.46 to 27.57 |
| Scaffold 2 | SJW-2C-239 | 33.07 | 21.97 to 55.43 |
| Scaffold 3 | SJW-2C-242 | 54.09 | 36.76 to 101.8 |

### Structure-activity relationships for the SJW-2C compounds

Analysis of the structures of the compounds from the hit expansion showed a number of insights. Compounds in scaffolds 1A and 1B (Tables S2 and S3) suggest that a cyclic alkyl, aryl, or aromatic moiety at position A_2_ would be active against the virus while an aliphatic alkyl chain would require a methyl group at the R_1_ position. Also, these compounds suggest that a methylene linker between the amide and A2 aryl group would be detrimental to its anti-viral activity. Further, based on compounds in Table 2, aromatic rings including phenyl, pyridine, thiophene, or furan groups at the A_2_ position result in active compounds. Further exploration of this chemical space would be needed to understand the SAR for the side chains at the A_2_ and R_1_ positions. Additional compounds for scaffolds 2 (Table S4) and 4 (Table S6) are needed to establish SAR. Compounds with scaffold 3 (Table S5) suggest that a methyl group at the 2-position of the furan ring as seen in SJW-2C-246 appears to be unfavorable to the function of that structure since it is absent in SJW-2C-245, SJW-2C-247 and SJW-2C-248, all of which have activity.

### SJW-2C-227 binds to recombinant EV-A71 2C protein *in vitro*

Due to limited quantities of other compounds, SJW-2C-227 was selected for subsequent assays. We used differential scanning fluorimetry (DSF) to study the direct binding of SJW-2C-227 to EV-A71 Δ2C^116-329^. Using this technique, an increase in the melting temperature (T_m_) of the target protein is observed if a compound binds to and stabilizes it. DSF has been used previously to show small molecule interactions with recombinant 2C^47,48^. An increase in the T_m_ was detected when SJW-2C-227 was incubated with Δ2C^116-329^ at a range from 8 µM to 40 µM (Figure 6). The shift in T_m_ indicated that the compound directly interacted with the Δ2C^116-329^ and that the inhibition of the ATPase function of Δ2C^116-329^ is likely a result of this direct interaction.

**Figure 6.**
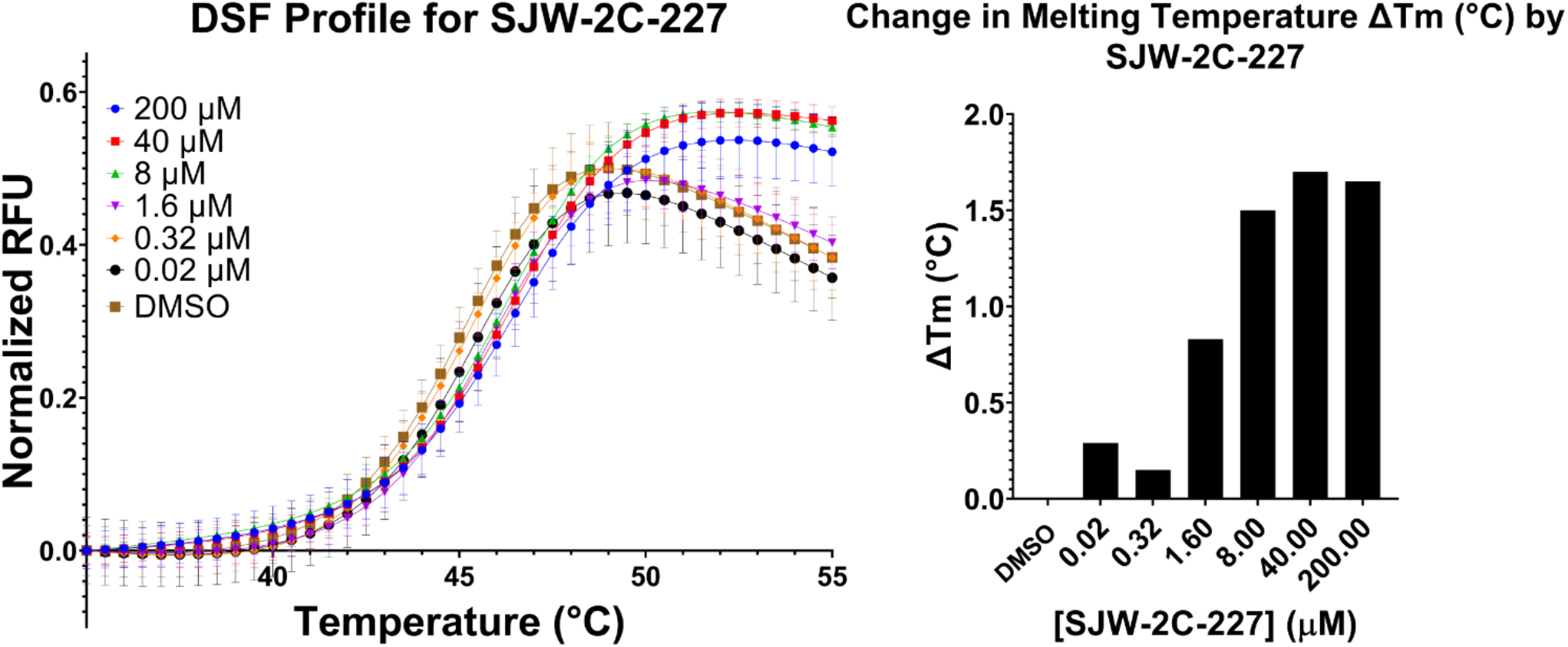
Differential scanning fluorimetry assay. Normalized relative fluorescence units (RFU) is plotted against the Temperature (°C) to give the melting profile of recombinant EV-A71 Δ2C^116-329^ in presence of increasing concentrations of the compound SJW-2C-227 (left panel). The reaction volume was 50 µL. Graphed mean and SD, n=3. The change in melting temperature when compared to the DMSO treated control (right panel).

### Escape mutants are located in the capsid proteins

We employed an escape mutant assay to provide insight into the binding site and mode of action of the compounds. Resistance to two compounds, SJW-2C-227, and its parent compound SJW-2C-69, was selected by serially passaging EV-A71 in the presence of increasing concentrations of the compounds. After 15 passages, viral genomes were extracted and sequenced. DMSO-treated virus served as a negative control and all the genome sequences from the drug-treated viruses were compared to the DMSO-treated EV-A71.

Passaging with the parent compound, SJW-2C-69, selected one mutation, T237N, in the VP1 capsid protein. To validate whether this mutation resulted in resistance, VP1-T237N was reverse engineered into the viral genome, and the mutant EV-A71-TN was generated. Comparative analysis using plaque assays showed that the mutant has similar infectivity to the wild-type EV-A71 (WT), producing plaques of similar size and shape (Figure S9). In the presence of SJW-2C-69, EV-A71-TN showed a ∼ 5-fold decrease in viral titer, while the wild-type EVA-71 showed a ∼100-fold decrease in the viral titer (Figure 7) indicating that the T237N mutation in VP1 offers partial resistance to SJW-2C-69. Serial passaging EV-A71 in the presence of SJW-2C-227 selected mutations in VP4 (K47E) and 3A (R34W).

**Figure 7.**
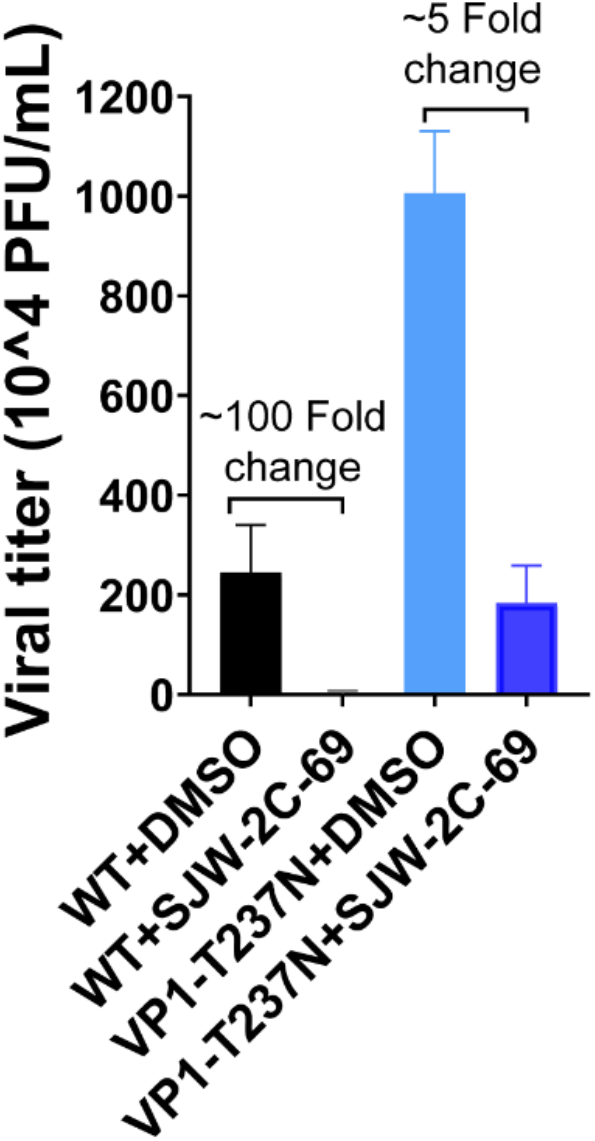
Validation of resistant escape mutant. A comparison of the viral titers for the DMSO treated vs the drug-treated, and the resulting fold change is shown as a bar graph. The wild type (WT) virus and the T237N mutant were generated from RNA as described previously. The final concentration of SJW-2C-69 used is 500 µM which is slightly less than 10x its IC_50_ (59.4 µM). Graphed mean and SD, n=3.

### SJW-2C-227 does not inhibit the EV-A71 replicon system

The DSF assay showed direct binding of the compounds to 2C, while the escape mutant assay suggested that the capsid proteins are involved in the mechanism of inhibition. To determine whether the compound was inhibiting the viral replication, we used an EV-A71 replicon system in which the P1 region, that encodes for the capsid proteins, is replaced with the sequence encoding for a fluorescent reporter protein whilst maintaining cleavage boundaries to ensure correct polyprotein processing. Replicon assays have previously been shown to be effective for the investigation of replication in real-time, independent of other aspects of the viral lifecycle such as entry, packaging, and egress^49–54^. *In vitro* transcribed WT and 3D^pol^-GNN (a replication-defective replicon harboring GDD>GNN mutation of the polymerase active site), EV-A71 replicon RNAs were transfected into HeLa cells treated with 1, 5, or 25 µM SJW-2C-227. The addition of the compound at all tested concentrations showed no statistically significant difference in the number of reporter positive cells when compared with the untreated control (Figure 8).

**Figure 8.**
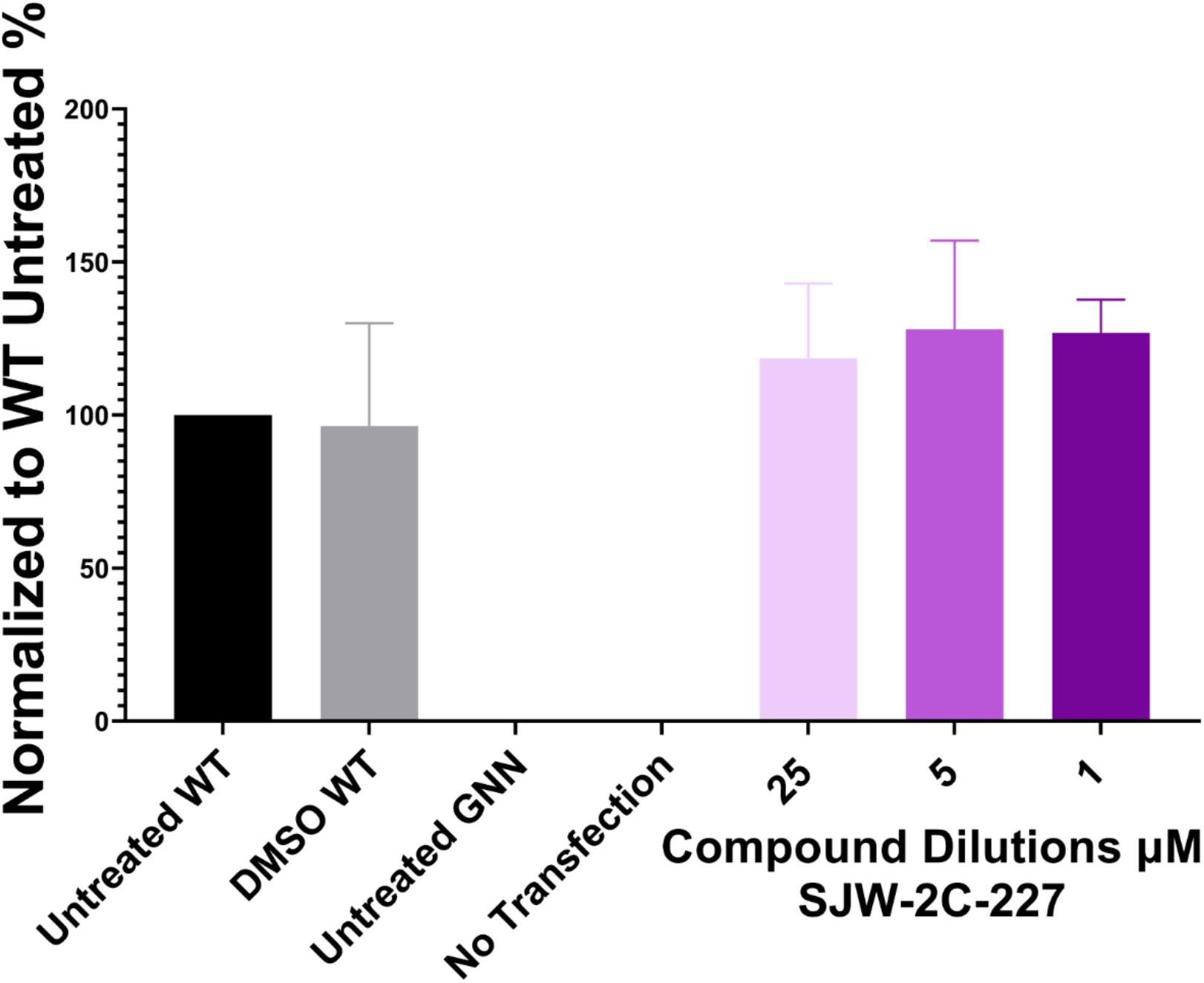
Effect of SJW-2C-227 on viral RNA replication using the EVA71 replicon system. Viral RNA levels were quantified upon transfecting Hela cells with the RNA generated from the EV-A71 replicon in the presence of either DMSO (vehicle control) or three different concentrations (25 µM, 5 µM, and 1 µM) of SJW-2C-227. In this replicon, the P1 capsid-encoding region is replaced with the sequence encoding for a fluorescent reporter protein. After 14 hours post-transfection, the fluorescence from the reporter protein was read. The untreated GNN (replication defective) and no transfection controls were used to remove the background fluorescence and the data were normalized to the untreated wild-type (WT) values. Graphed mean and SEM, n=3 in duplicate.

### Molecular Docking of SJW-2C-69 and SJW-2C-227 in the 2C:2C binding pocket

As the DSF and ATPase assays indicated a direct interaction between the two compounds and Δ2C^116-329^, we used *in silico* approaches to predict the interacting residues. We docked the parent compound, SJW-2C-69 and SJW-2C-227 in the 2C:2C binding pocket using Glide docking to identify potential intermolecular interactions with specific residues in this pocket^55–57^. Using the Induced Fit docking application with expanded sampling, 64 poses for SJW-2C-69 and 51 poses for SJW-2C-227 in the pocket were generated^58–60^. This program accounted for the flexibility of the ligand as well as the defined binding pocket. Figure 9 A and D show the top pose for SJW-2C-69 and SJW-2C-227 respectively and Figure 9 B and E show the residues forming the pocket interactions with the compounds. Applying the Interaction Fingerprints panel in Maestro to the top 9 poses of each ligand, the common residues interacting with the two compounds were identified (Figure 9 C and F)^61^.

**Figure 9.**
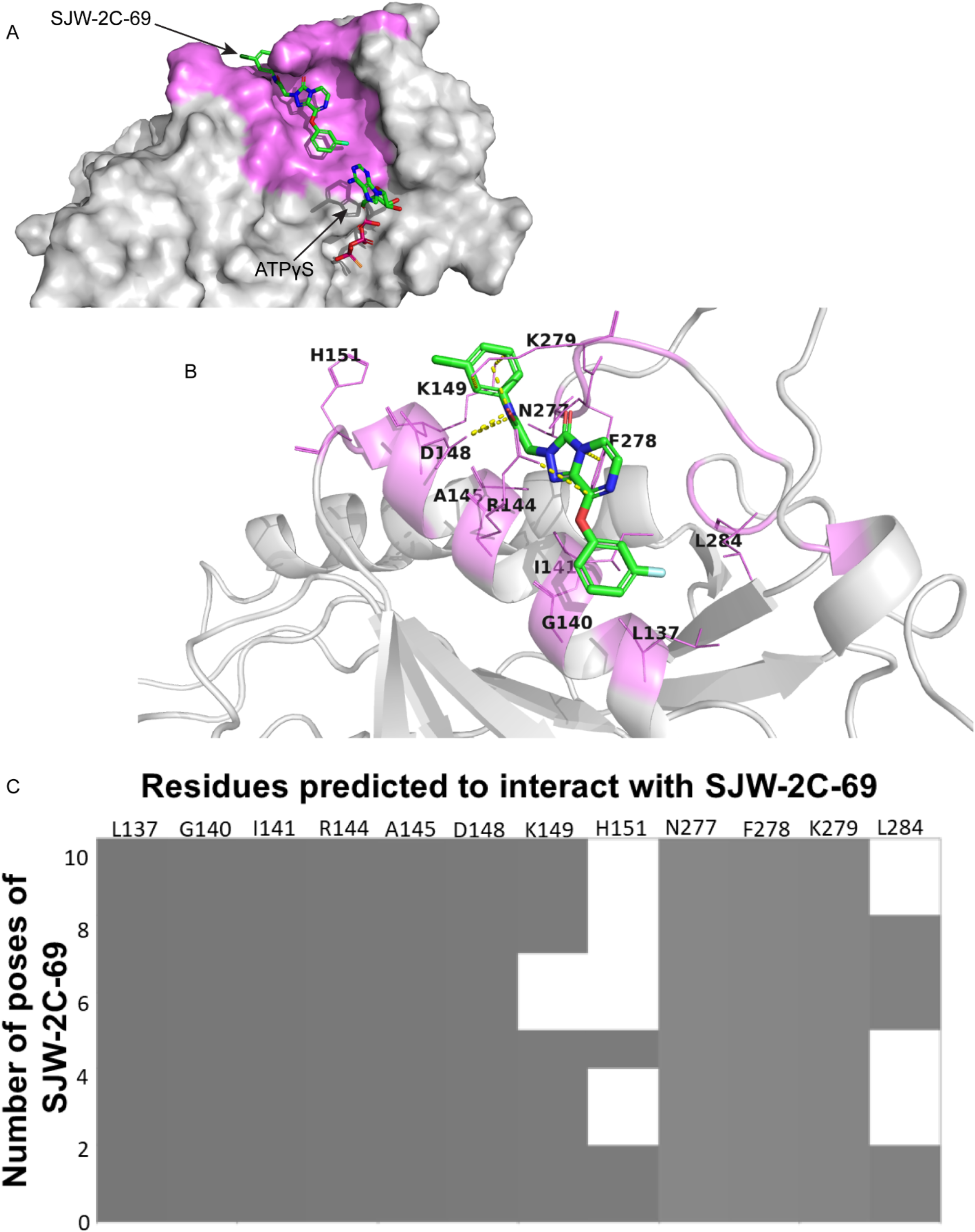
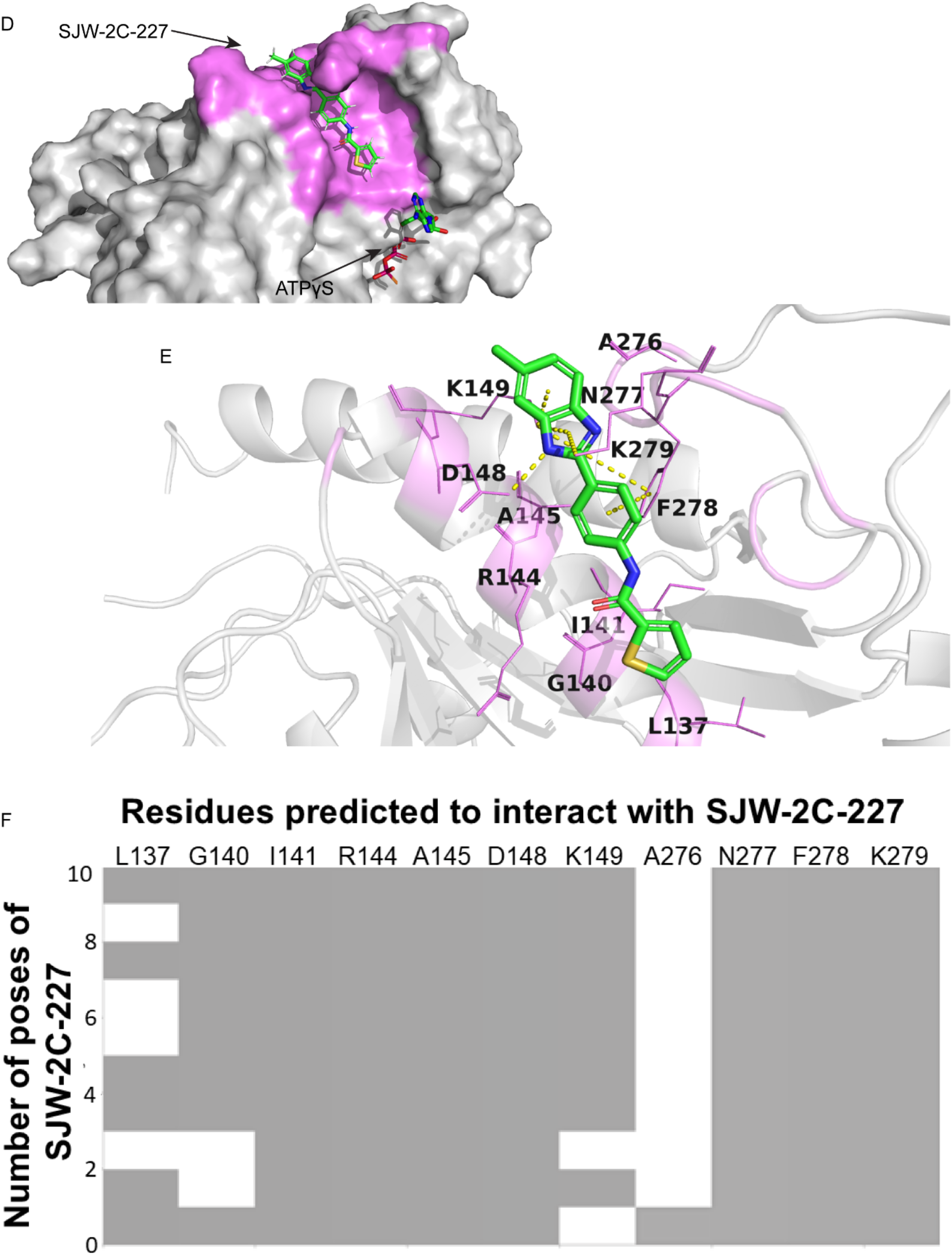
Molecular docking of SJW-2C-69 and SJW-2C-227 in the 2C:2C binding pocket to identify interacting residues. The two compounds, SJW-2C-69 and SJW-2C-227 (green stick diagram), were docked into the binding pocket (magenta) using the Glide docking application. The Induced Fit docking application was used to generate 64 poses for SJW-2C-69 and 51 poses for SJW-2C-234. Panels A and D show the top pose for the two ligands, SJW-2C-69 and SJW-2C-227, respectively in the pocket as a surface view. Panels B and E show the interacting residues in the pocket. Panels C and F show the common residues that interact with the compounds (gray box) when analyzing the top 10 poses using the Interaction Fingerprint application. The yellow dotted lines show the predicted pi-pi, pi-cation, and hydrogen bonds. All the applications are part of the Maestro suite from Schrödinger.

The top-ranked poses for the two compounds had a similar orientation and multiple contacts to residues in the pocket. The common residues interacting with SJW-2C-69 and SJW-2C-227 are L137, I141, R144, A145, D148, K149, A276, N277, F278, and K279. Residues L137, I141, A145, and A276 are predicted to contribute to hydrophobic interactions with the aromatic side chains of the compounds. R144 and K279 are predicted to form *π*-cation interactions while F278 forms *π* -*π* stacking interaction with the benzimidazole and the phenyl ring present in the two compounds. D148 is predicted to form hydrogen bonds with either the carboxyl group of the amide moiety in SJW-2C-69 or the benzimidazole ring of SJW-2C-227. L137 and I141 are immediately downstream of the Walker A motif. Further, in the absence of compound, R144 is suggested to form a salt bridge with E235 of the adjacent 2C subunit and help in the oligomerization of 2C. Wang *et al*. showed that the triple alanine mutation E148A/R149A/E150A in PV-1 2C resulted in increased virus temperature-sensitivity^18,62^. Although they confirmed that E150A alone resulted in temperature sensitivity, no further information was provided for the effects of the other two residues. N277 is conserved across the four representative EV species (Figure S1– N277 is shown as N279 due to multiple sequence alignment). K279 has been shown to be important at the encapsidation stage of the viral life cycle^17^. Several of the residues predicted to interact with SJW-2C-69 and SJW-2C-227 may have functional roles for the virus suggesting that mutating these residues might not produce a viable virus.

### SJW-2C-227 is active against EV-D68 and PV-1

A broad-spectrum anti-viral agent is highly desirable against enteroviruses. 2C is well conserved among enteroviruses, and we hypothesized that the compounds would have broad activity. To this end, we tested SJW-2C-227 against EV-D68, PV-1, and CV-B3. Results showed that SJW-2C-227 is highly potent against EV-D68 with an IC_50_ value of 0.85 µM and PV-1 strain with an IC_50_ of 1.7 µM, however no inhibition was observed against CV-B3 (Figure 10).

**Figure 10.**
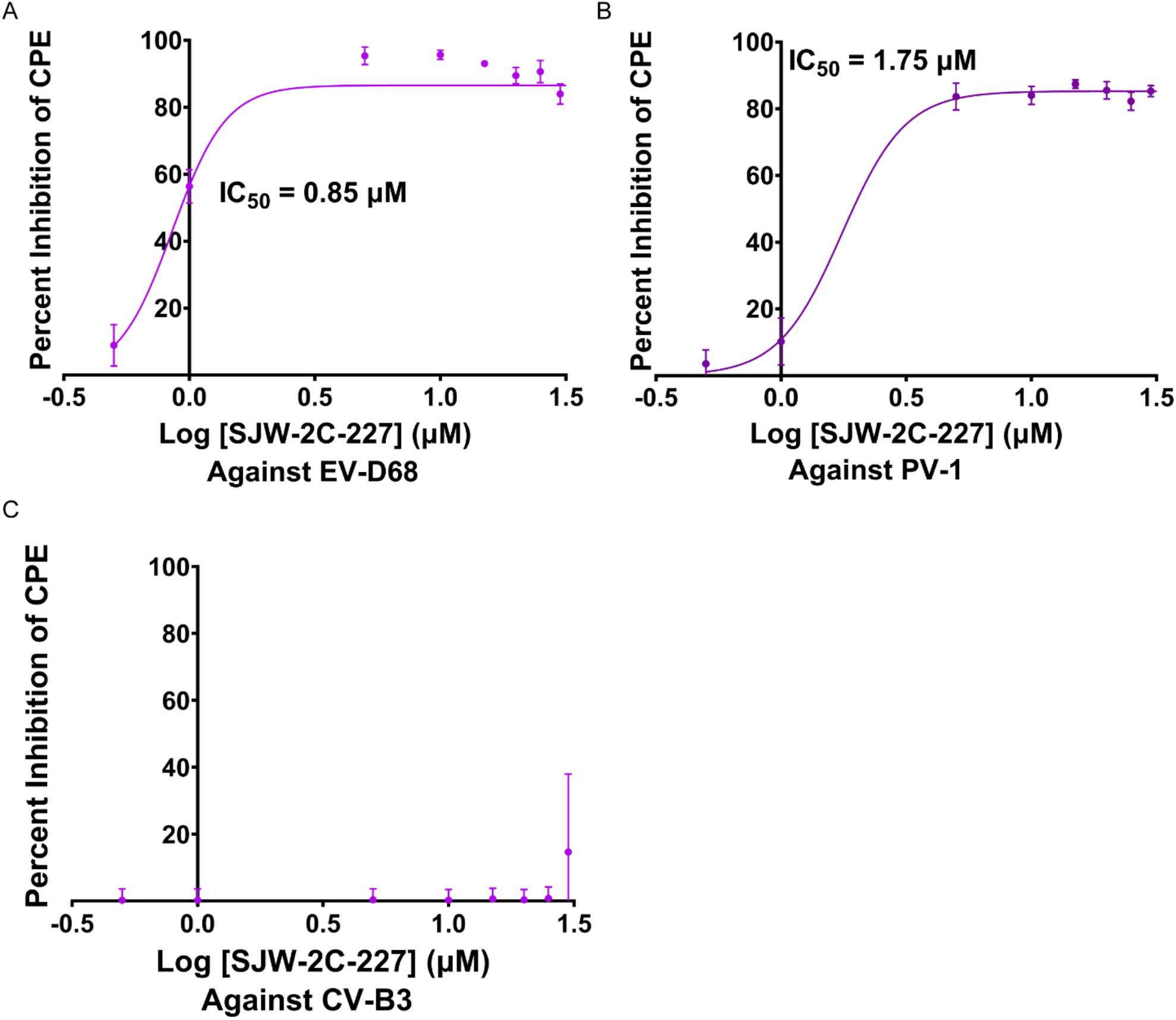
Broad spectrum activity of SJW-2C-227 against EV-D68 and PV-1. A dose-response curve was generated to determine the IC_50_ for the virus-induced CPE inhibition for SJW-2C-227 against EV-D68 (A), PV-1 (B) and CV-B3 (C). The assay was carried out in Vero cells for EV-D68 and PV-1 and HeLa cells for CV-B3. Graphed mean and SD, n=3.

We also used the Antiviral Program for Pandemics program (NIAID) to screen and validate the anti-viral results for SJW-2C-227 against a wider selection of viruses EV-A71, EV-D68, PV-1, CV-B3, eastern equine encephalitis virus, human echovirus-11, human echovirus-30, influenza A (H1N1), West Nile virus and Middle East respiratory syndrome. They used a different cytopathic effect assay which is desirable since it represents a robust validation of our anti-viral candidate. SJW-2C-227 was only active against EVA-71 and EV-D68 in this assay (Table 4) and showed no activity against the other viruses. SJW-2C-227 had an EC_50_ of 4.2 µM against EV-A71 and 0.39-0.52 µM against EV-D68. A secondary screen using a virus yield reduction assay was then carried out against EV-A71 and EV-D68. This is a two-step assay where first the virus is produced in cells containing the compounds and then the viral titer is measured. This assay confirmed the anti-viral activity of SJW-2C-227 with an EC_50_ of 1.7 µM against EV-A71 and 0.52 µM against EV-D68 (Table 5).

**Table 4.** Primary assay to test the antiviral activity of SJW-2C-227 through the Anti-viral Program for Pandemics from NIAID. EV, Enterovirus; CPE, Cytopathic effects; Tox, Toxicity; SI_50_ = CC_50_/EC_50_.

| PRIMARY ASSAY |  |  |  |  |  |
| --- | --- | --- | --- | --- | --- |
| Virus Name | Compound ID | Drug Assay Name | $EC_{50}$<br>$\mu M$ | $CC_{50}$<br>$\mu M$ | $SI_{50}$ |
| EV-71 | SJW-2C-227 | Visual (CPE/Tox) | > 32 | 32 | 0 |
|  | SJW-2C-227 | Neutral Red (CPE/Tox) | 4.7 | 42 | 8.9 |
| EV-68 | SJW-2C-227 | Visual (CPE/Tox) | 0.52 | 24 | 46 |
|  | SJW-2C-227 | Neutral Red (CPE/Tox) | 0.39 | 9 | 23 |

**Table 5.** Secondary assay to test the antiviral activity of SJW-2C-227 through the Anti-viral Program for Pandemics from NIAID. EV, Enterovirus; CPE, Cytopathic effects; Tox, Toxicity; VYR, Virus Yield Reduction; SI_50_ = CC_50_/EC_50_; SI_90_ = CC_50_/EC_90_

| **SECONDARY ASSAY** |  |  |  |  |  |  |  |
| --- | --- | --- | --- | --- | --- | --- | --- |
| **Virus Name** | **Compound ID** | **Drug Assay Name** | **EC<sub>50</sub>**<br>**μM** | **EC<sub>90</sub>**<br>**μM** | **CC<sub>50</sub>**<br>**μM** | **SI<sub>50</sub>** | **SI<sub>90</sub>** |
| EV-71 | SJW-2C-227 | Visual (VYR)/Neutral Red Tox |  | 3.2 | 27 |  | 7.8 |
|  | SJW-2C-227 | Neutral Red (CPE/Tox) | 1.7 |  | 25 | 15 |  |
| EV-68 | SJW-2C-227 | Visual (VYR)/Neutral Red Tox |  | 0.59 | 9.4 |  | 15.9 |
|  | SJW-2C-227 | Neutral Red (CPE/Tox) | 0.52 |  | 3.5 | 6.7 |  |

## Discussion

Enteroviruses are a major health burden globally but despite over 70 years of research efforts, no licensed small-molecule anti-EV therapeutics exist. 2C is highly conserved across the enteroviruses and is involved at multiple critical stages in the viral life cycle^27,31,33,39,63^. These properties make 2C a good target for broad-spectrum anti-viral therapeutics. Over the past four decades, several compounds have been identified that target 2C but to our knowledge, the compounds described in this paper are the first small molecules screened and selected to target the 2C:2C interaction pocket^43,46–48,51^.

Based on the X-ray structure of the EV-A71 Δ2C^116-329^, the C-terminal α6 helix of one 2C subunit protrudes into the 2C:2C binding pocket of the adjacent subunit and it was suggested that this interaction was important for the oligomerization of 2C^31^. Since 2C needs to oligomerize for ATPase activity, the inhibition of the ATPase activity served as the primary assay to determine whether the compounds were binding to the predicted 2C:2C interaction pocket. Inhibition of 2C ATPase function together with a dose-dependent increase in the T_m_ indicates that the compound binds directly to 2C. 2C is likely to exist in different conformational states during ATP hydrolysis, RNA binding, and helicase activities and it has been observed to exist in open and closed states^39^. SJW-2C-227 may preferentially bind to one state to partially inhibit ATPase function without completely inhibiting the ability of 2C to oligomerize and hydrolyze ATP. Even the most potent Δ2C^116-329^ ATPase inhibitor identified in this study, SJW-2C-219 (IC_50_ = 0.55 µM), had a maximum ATPase inhibition of ∼60-65 % at relatively high concentrations (Table 3). This suggests that the inhibition of the ATPase activity of 2C may not be the sole mechanism of action for its potent anti-viral activity observed in cells. Since 2C plays an important role in RNA replication^12–15,30^, the effect of viral RNA replication by this family of compounds was evaluated using a replicon system. It was observed that the compounds do not affect EV-A71 replication in the context of a replicon system (Figure S6).

Molecular docking studies predicted the orientation of the compound in the pocket and identified potential interacting residues. The pocket-forming residues are conserved amongst the enteroviruses (Figure S1) and so to escape inhibition by compounds it may be possible to select suppressor mutations in other proteins that are either directly affected by the compounds or in proteins that interact with 2C to avoid negative impact on its overall fitness. Consistent with this hypothesis, virus passaged in the presence of SJW-2C-69 and SJW-2C-227 evolved mutations in the capsid proteins VP1 (T237N) and VP4 (K47E), as well as the non-structural 3A protein (R34W). In a study performed by Wimmer and colleagues, enterovirus with an impaired encapsidation phenotype was rescued partially by a single mutation in either 2C (N252S) or in the capsid protein VP3 (E180G) but fully rescued by the double mutant^19^. This interaction between 2C and VP3 was also demonstrated biochemically using co-immunoprecipitation assays. Another study by Wang *et al*. showed that K279A/R280A in PV 2C produced encapsidation defective phenotypes^18^. Interestingly, K279 in EV-A71 forms a predicted pication bond with SJW-2C-227 (Figure 9 E). One of the rescue mutations from the Wang *et al*. study for this defect was T36I in the capsid protein VP1 but this mutant virus was temperature sensitive. For different clones, K41R in VP3 along with C323R in 2C or N203S in VP1 with C323R were also able to rescue the encapsidation defect^18^. These studies provide strong evidence that 2C interacts with the capsid proteins and plays a critical role in the morphogenesis of enteroviruses. Interestingly, residue 41 in EV-A71 VP3 is already an arginine and it is in proximity to the K47E mutation we observed in VP4 (Figure S7). Although these locations are based on the mature virus conformations, the region harboring these residues might be important for the virus encapsidation steps^64^. Wimmer and colleagues proposed a model where replication organelle-membrane-bound 2C interacts with the capsid pentamer through its interaction with VP3 and the pentamer then interacts with the VPg-linked viral genomic RNA that eventually leads to the assembly of progeny virus^19^. In a recent pre-print, using cryo-tomography on PV infected cells, Dahmane *et al*. observed that RNA loading of the capsids occurs while being tethered to the membrane^65^. Whether the tethering link is 2C still needs investigation. Further, Knox *et al*. observed that 2C and 3A along with VP1 and 3D, all co-localized to membrane clusters suggestive of replication organelles^66^. Recent studies have shown the role 3A plays in shuttling lipids between lipid droplets and replication organelles^67–71^. Yeast two-hybrid and mammalian two-hybrid systems have demonstrated interactions between 3A and 3AB with 2B, 2C, and 2BC ^14,72^. Interestingly, one of the escape mutations to SJW-2C-227 was in 3A – R34W. Whether this mutation imparts resistance still needs to be investigated.

Although the 2C:2C binding pocket is highly conserved, SJW-2C-227 was not active against CV-B3. Further, the escape mutant, VP1-T237N, was resistant to the parent compound, SJW-2C-69. Upon aligning the VP1 sequences from EV-A71, EV-D68, PV-1, and CV-B3, the equivalent position is either an isoleucine in EV-D68, a valine in PV-1, or a histidine in CV-B3 (Figure S8). Histidine at this location has been implicated to be responsible for the nuclear localization of CV-B3^73^. Whether this localization to the nucleus or nearby membranes is responsible for the phenotype needs further investigation. Since this histidine at this position is conserved among CV-B1-6^73^, determining whether SJW-2C-227 is effective against other CVs would be important for its development as a broad-spectrum antiviral agent.

Taken together, the results of this study suggest that the family of compounds may be inhibiting the interaction between 2C and the capsid proteins. Further, structural biology efforts are currently being conducted to determine a structure of the compound bound to 2C to aid the medicinal chemistry efforts to develop broad-spectrum antiviral agents with a novel chemistry against viruses that affect millions worldwide.

## Methods

### Cell lines, viruses, compounds, and chemicals

African Green Monkey kidney cells (Vero), H1 HeLa, and RD cells were purchased from ATCC. All the cell lines were cultured in Dulbecco’s modified Eagle’s medium (DMEM, Gibco, USA) supplemented with 10 % fetal bovine serum (FBS, Gibco, USA) and 1x Anti-Anti (Gibco, USA). All cell lines were maintained at 37 °C in 5 % CO_2_. EV-A71, PV-1, EV-D68, and CV-B3 were purchased from ATCC. EV-A71 and PV-1 were grown and amplified in Vero cells, EV-D68 was grown and amplified in RD cells and CVB3 was grown and amplified in H1 HeLa cells. The EV-A71 reverse genetics system and the EV-A71-T237N mutant were generated, grown and amplified as discussed previously^74^ but in Vero cells. Virus titers were determined by endpoint titration using plaque assay. The small molecule compounds were either supplied by Atomwise, Dr. Kyle Hadden (University of Connecticut), or purchased from Hit2Lead. Guanidinium hydrochloride, Dibucaine, and Fluoxetine were purchased from Sigma-Aldrich (USA). For recombinant protein production, the *E. coli* was grown in 2x YT medium. The 2x YT medium was prepared by mixing 16 g of Tryptone (Sigma-Aldrich, USA), 10 g of Yeast extract (Sigma-Aldrich, USA), 5 g of sodium chloride (Fisher Scientific, USA), and 2 g of glucose (Fisher Scientific, USA) in 1 L of de-ionized water and then sterilized by autoclaving for 30 minutes at 15 psi (1.05 kg/cm^2^) on liquid cycle.

### *In silico* screening

The available 2C structure (PDB: 5GRB) is a 2.8 Å crystal structure in which six 2C molecules fill the asymmetric unit^31^. The oligomerization of 2C has been observed to be important for its ATPase function, but it has been unclear whether the crystal contacts observed in the available structure are truly representative of the solution structure conformation. On comparing Chains, A – F of the PDB structure, there were six very different interfaces observed in the asymmetric unit, and only Chain F showed ATP binding in a reasonable orientation. Guan *et al*. built a model of the 2C hexamer but did not share the coordinates and suggested that it was very approximate^31^. Thus, chain F binding pocket was used as the protein receptor for virtual screening, with residues 323-329 of chain A serving as the prototype cognate binding ligand. The residues of the receptor pocket include L137, G140, I141, I142, R144, A145, D148, A267, K268, L269, N277, F278, K279, R280, C281, S282, L284, V285. Before screening, the ATP ligand, solvent, and ions were removed. A curated molecular library of several million compounds was screened in silico using the AtomNet® technology^35^. Top scoring compounds were clustered and subsequently filtered for favorable properties, for example, solubility, known toxicity, etc., to arrive at a final subset of 77 deliverable compounds.

### 2C viral protein purification

EV-A71 Δ2C (amino acids 116-329) was designed with an N-terminal MBP tag and a C-terminal Hexa-His tag. The DNA fragment encoding the protein was codon optimized for *Escherichia coli* (*E. coli)* protein expression and synthesized synthetically by Genscript (New Jersey, USA) before being cloned into the pMALc5x vector. The protein was produced in the *E. coli* Rosetta™ 2(DE3) pLysS cells (Millipore Sigma, USA). The cells were grown in the 2x YT media containing 100 μg/mL of ampicillin and 50 μg/mL of chloramphenicol at 37 °C until O.D. reached 0.6 after which the protein production was induced with 0.25 mM IPTG and the cells were grown for 18 hours at 20 °C. The cells were harvested by centrifugation at 8000xg for 10 minutes and frozen at -80 °C. The frozen pellets were resuspended in the lysis buffer (20 mM Bis-Tris pH 7.2, 200 mM NaCl, Roche’s complete Protease inhibitors EDTA-free, 5 mM TCEP, 1 % Triton X-100 and small quantities of DNase and RNase). The resuspended cells were lysed using the One-Shot Cell Disrupter (Constant Systems, UK) following which the lysate was incubated for 30 minutes at room temperature. The lysate was centrifuged at 25000 xg for 30 minutes to remove cell debris and the supernatant was filtered through a 0.22μm membrane filter (Millipore Sigma, USA). Imidazole equivalent to 50 mM final concentration was added to the filtrate containing the MBP-Δ2C-(His)_6_ protein. The protein was first purified using the Ni-NTA affinity chromatography (HisTrap Cytiva, USA) and then through size exclusion chromatography using the Superdex S200 10/300 column (Cytiva, USA). The affinity chromatography buffer A contained 20 mM Bis-Tris pH 7.2, 200 mM NaCl, and 0.1 % Triton X-100 while the elution buffer contained 500 mM imidazole in addition to buffer A components. SEC buffer contained 20 mM Bis-Tris pH 7.2 and 200 mM NaCl.

### 2C ATPase assay

The ATPase activity of the MBP-Δ2C-(His)_6_ was confirmed using the colorimetric malachite green assay. This assay was performed in 384-well plates. Each 50 μL reaction contained 2 μM of Δ2C protein, a fixed concentration of the compound or DMSO, 0.5 μM ATP, 5 mM MgCl_2,_ and 20 mM Bis-Tris pH 7.2 and was incubated at room temperature for 18 hours before adding 100 μL of the color reagent: 0.045 % malachite green (Fisher Scientific), 34 mM ammonium molybdate (Fisher Scientific) in 4 N HCl and 0.02 % Tween-20 (Fisher Scientific). The plate was then incubated at room temperature for 15 minutes before reading the absorbance at 660 nm. The inhibitory effects of the small molecule compounds were first measured at 50 μM after which a dose-response curve was generated for the hits to calculate their IC_50_ values. The final DMSO concentration was kept constant at 10 % after ensuring that the ATPase activity of 2C was not inhibited by 10 % DMSO. GnHCl and the racemic mixture of Fluoxetine were tested as positive controls and so the appropriate IC_50_ value was used as a reference.

For the dose-response curves, the percent inhibition of ATPase activity was calculated according to the following formula and plotted against the drug concentration:

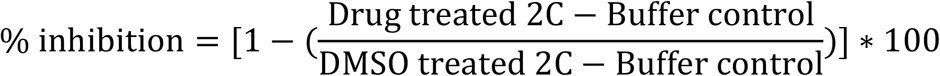

Curve fitting was carried out in GraphPad Prism using non-linear regression analysis and the IC_50_ was calculated using the log(inhibitor) vs. response - variable slope model. The assay was performed three times from which the 95 % confidence interval range for IC_50_ was calculated using profile likelihood asymmetrical confidence intervals. The bottom constraint was set as zero and the top constraint was set as 100 only in cases where a plateau was not observed at higher compound concentrations.

### Differential Scanning Fluorimetry (DSF)

The binding of the compounds to 2C was assessed by studying the thermal stability of 2C by DSF using Sypro Orange (ThermoFisher Scientific). This was monitored using the Bio-Rad CFX96 Touch Real-Time PCR Detection System using the FRET channel. Each 50 μL reaction contained 2 μM of Δ2C^116-329^ protein and a fixed concentration of the compound that was incubated at room temperature for 30 minutes. Following incubation, Sypro Orange at 5x concentration was added to the reaction and the plate was read for fluorescence. The final DMSO concentration was 4 % before the addition of Sypro Orange. The compounds were tested at six different concentrations (200 μM, 40 μM, 8 μM, 1.6 μM, 0.32 μM and 0.02 μM) to study the dose-dependent shift in the melting temperature (T_m_) of 2C upon interaction with the compound.

### Molecular docking

The molecular docking studies were carried out using the Schrödinger suite (v. 2022-1). First, the compound structures were downloaded as a .sdf from PubChem. In the Maestro application, the ligands were prepped using the LigPrep application. The 2C protein structure (PDB: 5GRB, Chain F) was loaded and prepped using the default settings in the Protein Preparation wizard including H-bond optimization and energy minimizations. Docking experiments were performed using the Induced Fit Docking (extended sampling) application in Maestro. Once the docked poses were generated, the top 10 poses were selected to generate the common residue interaction profile using the Interaction Fingerprint application.

### Similarity study

All the computational studies were performed using either Maestro, the graphical user interface for Schrodinger Suite (2021), or Canvas. The three-dimensional structure of each molecule was generated using the ‘build’ module within Schrodinger. Hashed binary fingerprints of each compound were generated by linear fragments with the ring closure method. Using the binary fingerprints as our geometric atomic descriptor, the similarity study was carried out using the Tanimoto association coefficient, which gives the similarity values on a scale from 0 to 1. The Tanimoto distance metric is a normalized measure of the similarity in descriptor space between a series of test compounds and a probe molecule. Similarities lie between one and zero with a value of one indicating identical molecules and a value of zero indicating completely dissimilar molecules. Further hierarchical clustering was performed using the binary fingerprints generated for the compounds.

The Tanimoto distance metrics were calculated as follows:

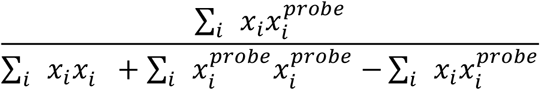

In this equation, *x*_*i*_ = values of data point i in test molecules, *x*_*iprobe*_ = values of data point *i* in probe molecule 1, *x*_*i*_*x*_*iprobe*_ = values of common data point *i* in both test and probe molecule.

### Multiplicity of Infection (MOI) screening

Multiplicity of infection (MOI) is defined as the number of plaque-forming units (PFU) of the virus present per host cell. We screened for an MOI that would result in complete cell death after 3 days of infection. This MOI was first determined for each of the viruses in their respective cell system. For example, for EV-A71, Vero cells were used. For this assay, the Cell Titer Glo One solution kit from Promega was used. Vero cells were seeded in 96-well white plates at a density of 25,000 cells per well and incubated overnight at 37 °C in a 5 % CO_2_ incubator. The next day, multiple dilutions of the viral stock with a known titer (PFU/mL) were prepared using serum-free DMEM (supplemented with 1x Anti-Anti) such that a range of MOIs were screened. The highest MOI tested was 10 and a 2x dilution series was prepared. 200 μL of the respective dilution of the virus was added per well. The virus was adsorbed for 2 hours in the incubator. After 2 hours, the virus was removed and 250 μL growth media (DMEM + 10 % FBS + 1x Anti-Anti) was added to each well. Control wells containing cells without virus were also included. The plate was incubated for 3 days. After 3 days, the Cell Titer Glo One solution kit was used to measure the cell viability. 150 μL of the media from each well was discarded and 100 μL of the Cell Titer Glo reagent was added to each well. The plate was carefully shaken for 2 minutes and then incubated at room temperature in a biosafety cabinet for 10 minutes before reading the luminescence. The relative luminescence units (RLU) was then plotted against the MOI and the minimum MOI at which all the cells were dead (lowest luminescence) was noted.

### CPE inhibition assay

Anti-viral activity of the compounds on EV-A71, EV-D68, poliovirus, and coxsackievirus B3 was carried out using a cytopathic effect (CPE) inhibition assay. For this assay, the Cell Titer Glo One solution kit from Promega was used. Vero cells were seeded in 96-well white plates at a density of 25,000 cells per well and were incubated overnight at 37 °C in a 5 % CO2 incubator. The next day, the cells were infected with EV-A71 at an MOI of 1 and the virus was left to adsorb for 2 hours in the incubator at 37 °C and 5 % CO_2_. After 2 hours, the virus supernatant was discarded, and the compound was added at a fixed concentration of either 50 μM or 25 μM in fresh media, and the plate was incubated for 3 days at 37 °C and 5 % CO_2_. The final DMSO concentration in each well was kept constant at 1 %. For dose-response curves, instead of one concentration of the compound, a range of concentrations from 0.1 μM to 50 μM, were used. After 3 days, the Cell Titer Glo One solution kit was used to measure the cell viability. 150 μL of the media from each well was discarded and 100 μL of the Cell Titer Glo reagent was added to each well. The plate was carefully shaken for 2 minutes and then incubated at room temperature in the biosafety cabinet for 10 minutes before reading the luminescence.

For the IC_50_ calculations, the percent inhibition of cytopathic effect (CPE) was plotted against the drug concentration. Percent CPE was calculated using the following formula:

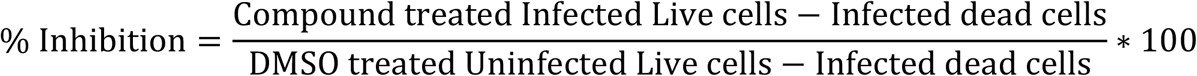

Curve fitting was carried out in GraphPad Prism using non-linear regression analysis and the IC_50_ was calculated using the log(inhibitor) vs. response - variable slope model. Further, the curve fitting was carried out with a bottom constraint of 0 while a top constraint of 100 was added to plots that did not reach a plateau at higher compound concentrations. The assay was performed three times from which the 95 % confidence interval range for IC_50_ was calculated using profile likelihood asymmetrical confidence intervals.

### Cytotoxicity Assay

The cellular toxicity of the compounds was tested using the cytotoxicity assay and a dose-response assay was performed to determine the CC_50_ for each of the compounds. For this assay, the Cell Titer Glo One solution kit from Promega was used. Vero cells were seeded in 96-well white plates at a density of 25,000 cells per well and was incubated overnight at 37 °C in a 5 % CO_2_ incubator. The next day, serum-free media was added to the cells and the plate was incubated for 2 hours in the incubator at 37 °C and 5 % CO_2_. After 2 hours, the media was discarded, and the compound was added, and the plate was incubated for 3 days at 37 °C and 5 % CO_2_. The compound was dissolved in DMSO and the required concentration was prepared in growth media. Further, the final DMSO concentration in each well was kept constant at 1 %. For dose-response curves, a range of concentrations from 0.5 μM to 1000 μM were used depending upon the availability of the compounds. After 3 days, the Cell Titer Glo One solution kit was used to measure the cell viability. 150 μL of the media from each well was discarded and 100 μL of the Cell Titer Glo reagent was added to each well. The plate was carefully shaken for 2 minutes and then incubated at room temperature in the biosafety cabinet for 10 minutes before reading the luminescence.

The CC_50_ values were calculated by plotting percent viability against the drug concentration.

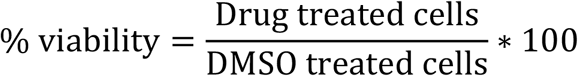

Curve fitting was carried out in GraphPad Prism using non-linear regression analysis and the IC_50_ was calculated using the log(inhibitor) vs. response - variable slope model. Further, the curve fitting was carried out with a bottom constraint of 0 in cases where the compound was not cytotoxic at higher concentrations and percent viability did not reach 0 %. The assay was performed three times.

### EV-A71 Replicon Assay

HeLa cells were maintained using Dulbecco’s modified Eagle’s medium (DMEM) containing 10 % (v/v) fetal bovine serum (Sigma-Aldrich), and 1 % Glutamax (ThermoFisher Scientific), and penicillin-streptomycin (Sigma-Aldrich) (termed complete DMEM, cDMEM). EV71 replicon plasmid was linearized using *Xho*I (NEB) as previously described^75^ before *in vitro* transcription using the T7 RiboMAX system (Promega) following the manufacturer protocol but with all volumes and concentrations reduced by a factor of two. RNA transcripts were recovered using the RNA Clean and Concentrator-25 kit (Zymo Research), with concentrations determined by NanoDrop (ThermoFisher Scientific) and RNA integrity determined by MOPS-formaldehyde denaturing gel electrophoresis. HeLa cells were seeded 16 hours before RNA transfection at a density of 10,000 cells per well of a 96-well plate. Wild-type and 3D-GNN (replication-defective mutant) RNAs were prepared by diluting 160 ng/well RNA and 0.48 µL/well Lipofectamine 2000 (ThermoFisher Scientific) in opti-MEM™ (ThermoFisher Scientific). Following a 10-minute incubation, RNA and lipofectamine mixes were combined and incubated for a further 20 minutes. The transfection mix was added to cDMEM without phenol-red indicator, alongside chemical compounds where applicable, to a final volume of 100 µL/well. Cells were washed 1X with PBS before the addition of RNA transfection/cDMEM mixture to wells in triplicate. Four images of each well were taken hourly using an IncuCyte S3 Live Cell Imaging System (Sartorius) to visualize fluorescent reporter protein expression. Fluorescent cells were determined using the inbuilt IncuCyte analysis package using surface fit segmentation and filters to emit any signal with a threshold value (GCU; an arbitrary green fluorescent unit) <10 and surface area < 50 µm^2^; values that were determined using no transfection and 3D-GNN controls to remove background fluorescence. The average number of positive cells per condition 14 hours post-transfection across triplicates was compared to untreated wild-type transfected wells. Averages of three independent repeats were plotted for each sample and statistical analysis was performed using a two-way ANOVA unless otherwise indicated.

### Escape Mutant Assay

Escape mutant assay involves selecting resistant viruses to specific compounds to determine their mechanism of action. To select escape mutants for SJW-2C-69, Vero cells were seeded at a density of 25,000 cells per well of a 96-well plate and incubated overnight at 37 °C in a 5 % CO_2_ incubator. The next day, the cells were infected with EV-A71 at an MOI of 5 and the virus was left to adsorb for 2 hours in the incubator at 37 °C and 5 % CO_2_. After 2 hours, the virus was discarded, and the compound was added at a fixed concentration of 0.5x IC_50_ value (30 µM) and the plate was incubated for 3 days at 37 °C and 5 % CO_2_. After 3 days, the cells were checked for cytopathic effects (CPE) using a light microscope. CPE indicated successful viral infection and the supernatant media contained the progeny virus. This virus was termed generation 1 (Gen 1). The virus from Gen 1 was used to infect a freshly seeded plate of Vero cells. Following the 2-hour adsorption, the virus was removed and saved. Fresh media containing 0.5x IC_50_ value (30 µM) of SJW-2C-69 was added to the cells and incubated for 3 days at 37 °C and 5 % CO_2_. The supernatant media containing the virus was then used to infect a new batch of Vero cells and the process is repeated until 15 generations of the virus were produced including five generations with 0.5x IC_50_ value (30 µM), followed by five generations with 1x IC_50_ (60 µM) and finally, five generations with 2x IC_50_ (120 µM). Escape mutants for SJW-2C-227 were extracted similarly with a few modifications. The 15 generations were produced such that there were three generations at five different drug concentrations – 0.5x IC_50_, 1x IC_50_, 2x IC_50_, 4x IC_50,_ and 10x IC_50_. A similar approach was followed to generate the DMSO-treated virus as well to compare the mutations. The DMSO concentration was kept constant at 1 % throughout the 15 generations. The viral genome of Gen 15 was isolated using TRIzol extraction (Sigma-Aldrich). The genome was then sequenced using next-generation sequencing.

### Total RNA QC

Total RNA was quantified, and purity ratios were determined for each sample using the NanoDrop 2000 spectrophotometer (Thermo Fisher Scientific, Waltham, MA, USA). To assess RNA quality, total RNA was analyzed on the Agilent TapeStation 4200 (Agilent Technologies, Santa Clara, CA, USA) using the RNA High Sensitivity assay following the manufacturer’s protocol. Ribosomal Integrity Numbers (RINe) were recorded for each sample.

### Illumina total RNA library preparation and sequencing

Total RNA samples (250ng of Qubit quantified total RNA input) were prepared for library preparation using the Illumina Stranded Total RNA Gold library preparation kit (Illumina, San Diego, CA) following the manufacturer’s protocol. Libraries were validated for length and adapter dimer removal using the Agilent TapeStation 4200 D1000 High Sensitivity assay (Agilent Technologies, Santa Clara, CA, USA) then quantified and normalized using the dsDNA High Sensitivity Assay for Qubit 3.0 (Life Technologies, Carlsbad, CA, USA).

Sample libraries were prepared for Illumina sequencing on the NovaSeq 6000 by denaturing and diluting the libraries per the manufacturer’s protocol (Illumina, San Diego, CA, USA). All samples were combined into one sequencing pool, proportioned according to the expected number of reads, and run as one sample pool. Target read depth was achieved per sample with paired-end 150bp reads.

### Plaque assay for validation of the resistant escape mutant

The EV-A71 reverse genetics system and the EV-A71-T237N mutant were generated, grown and amplified as discussed previously^74^ with the exception that the virus was grown and amplified in Vero cells. To compare the resistance of the escape mutant with wild-type, sufficient wild-type (WT) and T237N mutant virus stock that would result in complete cell death after 3 days post-infection was used as the starting virus concentration. Six serial dilutions were carried out of this virus and used in plaque assay as described here to determine the viral titer in the presence of 500 µM of SJW-2C-69. Vero cells, cultured in growth media (DMEM + 10 % FBS + 1x Anti-Anti), were seeded at a density of 500,000 cells/well in a clear 6-well plate and incubated overnight at 37 °C in a 5 % CO_2_ incubator. The next day, six 10-fold serial dilution of the virus was prepared in serum-free DMEM (with 1x Anti-Anti). 500 μL of the dilution was added to the cells and the plate was incubated at 37 °C in a 5 % CO_2_ incubator. The plate was rocked gently every 20 minutes to ensure even coverage and prevent the cellular monolayer from drying. After 2 hours of adsorption, the virus was removed, and the cells were washed with phosphate-buffered saline (PBS). 2 mL of the liquid overlay was added to each well and the plate was incubated at 37 °C in a 5 % CO_2_ incubator for 3 days. The liquid overlay was prepared by mixing 1:1 2.4 % Avicel (DuPont) in PBS and growth media so that the final concentration of Avicel was 1.2 %. When testing the effect of SJW-2C-69 on the viral titer, the compound was added to the liquid overlay at a final concentration of 500 µM which is slightly lower than 10x its IC_50_ (59.4 µM). After 3 days the cells were fixed and stained. To fix the cells, 1 mL of 4 % formaldehyde was added to each well and incubated at room temperature in a biosafety cabinet for 30 minutes. Then the formaldehyde-Avicel overlay was poured out and washed with de-ionized water. 1 mL of 1 % crystal violet staining solution was then added to each well. After 1 hour the stain was discarded and the wells were rinsed in water first and then in 0.5 % bleach. The plates were dried in a biosafety cabinet overnight. Once the plate was dry, the viral titer was determined by the following formula. –

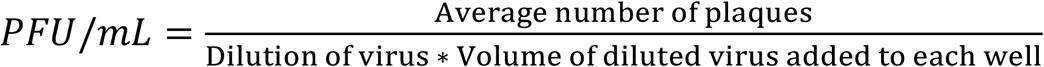

Example - If there were 32 plaques in the well which was infected with 500 mL of the 10^−5^ virus dilution, then the viral titer would be 64 × 10^−5^ PFU/mL [32 plaques / (10^−5^ * 0.5 mL)].

## Acknowledgments

We wish to thank Dr. Bo Reese at the University of Connecticut Center for Genome Innovation for help with the Next-generation Sequencing. We thank the Connecticut PITCH Program in Innovative Therapeutics for Connecticut’s Health for funding. We gratefully acknowledge support from The UK Medical Research Council MR/P022626/1 (NJK, NJS, DJR), UK Biotechnology and Biological Sciences Research Council BB/T015748/1(SJD, NJS and DJR) and the NIH R01 AI 169457-0 (NJK, NJS, DRJ). In addition, NJK received a fellowship from the Wellcome trust ISSF (204825/Z/16/Z)

## Notes and conflict of interest

The authors declare the following competing financial interest(s): R.K. and S.J.W. are inventors of a pending patent that claimed the use of the described compounds as potential broad-spectrum antivirals against Picornaviruses.

## References

(1) Nikonov, O. S.; Chernykh, E. S.; Garber, M. B.; Nikonova, E. Y. Enteroviruses: Classification, Diseases They Cause, and Approaches to Development of Antiviral Drugs. Biochem. Biokhimiia 2017, 82 (13), 1615–1631. https://doi.org/10.1134/S0006297917130041.

(2) Tapparel, C.; Siegrist, F.; Petty, T. J.; Kaiser, L. Picornavirus and Enterovirus Diversity with Associated Human Diseases. Infect. Genet. Evol. J. Mol. Epidemiol. Evol. Genet. Infect. Dis. 2013, 14, 282–293. https://doi.org/10.1016/j.meegid.2012.10.016.

(3) Chang, L.-Y.; Lin, H.-Y.; Gau, S. S.-F.; Lu, C.-Y.; Hsia, S.-H.; Huang, Y.-C.; Huang, L.-M.; Lin, T.-Y. Enterovirus A71 Neurologic Complications and Long-Term Sequelae. J. Biomed. Sci. 2019, 26 (1), 57. https://doi.org/10.1186/s12929-019-0552-7.

(4) Puenpa, J.; Wanlapakorn, N.; Vongpunsawad, S.; Poovorawan, Y. The History of Enterovirus A71 Outbreaks and Molecular Epidemiology in the Asia-Pacific Region. J. Biomed. Sci. 2019, 26 (1), 75. https://doi.org/10.1186/s12929-019-0573-2.

(5) Pons-Salort, M.; Parker, E. P. K.; Grassly, N. C. The Epidemiology of Non-Polio Enteroviruses: Recent Advances and Outstanding Questions. Curr. Opin. Infect. Dis. 2015, 28 (5), 479–487. https://doi.org/10.1097/QCO.0000000000000187.

(6) Schubert, R. D.; Hawes, I. A.; Ramachandran, P. S.; Ramesh, A.; Crawford, E. D.; Pak, J. E.; Wu, W.; Cheung, C. K.; O’Donovan, B. D.; Tato, C. M.; Lyden, A.; Tan, M.; Sit, R.; Sowa, G. M.; Sample, H. A.; Zorn, K. C.; Banerji, D.; Khan, L. M.; Bove, R.; Hauser, S. L.; Gelfand, A. A.; Johnson-Kerner, B. L.; Nash, K.; Krishnamoorthy, K. S.; Chitnis, T.; Ding, J. Z.; McMillan, H. J.; Chiu, C. Y.; Briggs, B.; Glaser, C. A.; Yen, C.; Chu, V.; Wadford, D. A.; Dominguez, S. R.; Ng, T. F. F.; Marine, R. L.; Lopez, A. S.; Nix, W. A.; Soldatos, A.; Gorman, M. P.; Benson, L.; Messacar, K.; Konopka-Anstadt, J. L.; Oberste, M. S.; DeRisi, J. L.; Wilson, M. R. Pan-Viral Serology Implicates Enteroviruses in Acute Flaccid Myelitis. Nat. Med. 2019, 25 (11), 1748–1752. https://doi.org/10.1038/s41591-019-0613-1.

(7) Park, S. W.; Pons-Salort, M.; Messacar, K.; Cook, C.; Meyers, L.; Farrar, J.; Grenfell, B. T. Epidemiological Dynamics of Enterovirus D68 in the United States and Implications for Acute Flaccid Myelitis. Sci. Transl. Med. 2021, 13 (584), eabd2400. https://doi.org/10.1126/scitranslmed.abd2400.

(8) Vogt, M. R.; Wright, P. F.; Hickey, W. F.; De Buysscher, T.; Boyd, K. L.; Crowe, J. E. Enterovirus D68 in the Anterior Horn Cells of a Child with Acute Flaccid Myelitis. N. Engl. J. Med. 2022, 386 (21), 2059–2060. https://doi.org/10.1056/NEJMc2118155.

(9) CDC. AFM Cases and Outbreaks. Centers for Disease Control and Prevention. https://www.cdc.gov/acute-flaccid-myelitis/cases-in-us.html (accessed 2022-06-02).

(10) Konopka-Anstadt, J. L.; Campagnoli, R.; Vincent, A.; Shaw, J.; Wei, L.; Wynn, N. T.; Smithee, S. E.; Bujaki, E.; Te Yeh, M.; Laassri, M.; Zagorodnyaya, T.; Weiner, A. J.; Chumakov, K.; Andino, R.; Macadam, A.; Kew, O.; Burns, C. C. Development of a New Oral Poliovirus Vaccine for the Eradication End Game Using Codon Deoptimization. Npj Vaccines 2020, 5 (1), 1–9. https://doi.org/10.1038/s41541-020-0176-7.

(11) Famulare, M.; Chang, S.; Iber, J.; Zhao, K.; Adeniji, J. A.; Bukbuk, D.; Baba, M.; Behrend, M.; Burns, C. C.; Oberste, M. S. Sabin Vaccine Reversion in the Field: A Comprehensive Analysis of Sabin-Like Poliovirus Isolates in Nigeria. J. Virol. 2015, 90 (1), 317–331. https://doi.org/10.1128/JVI.01532-15.

(12) Li, J. P.; Baltimore, D. Isolation of Poliovirus 2C Mutants Defective in Viral RNA Synthesis. J. Virol. 1988, 62 (11), 4016–4021. https://doi.org/10.1128/JVI.62.11.4016-4021.1988.

(13) Rieder, E.; Paul, A. V.; Kim, D. W.; van Boom, J. H.; Wimmer, E. Genetic and Biochemical Studies of Poliovirus Cis-Acting Replication Element Cre in Relation to VPg Uridylylation. J. Virol. 2000, 74 (22), 10371–10380. https://doi.org/10.1128/jvi.74.22.10371-10380.2000.

(14) Teterina, N. L.; Levenson, E.; Rinaudo, M. S.; Egger, D.; Bienz, K.; Gorbalenya, A. E.; Ehrenfeld, E. Evidence for Functional Protein Interactions Required for Poliovirus RNA Replication. J. Virol. 2006, 80 (11), 5327–5337. https://doi.org/10.1128/JVI.02684-05.

(15) Tang, W.-F.; Yang, S.-Y.; Wu, B.-W.; Jheng, J.-R.; Chen, Y.-L.; Shih, C.-H.; Lin, K.-H.; Lai, H.-C.; Tang, P.; Horng, J.-T. Reticulon 3 Binds the 2C Protein of Enterovirus 71 and Is Required for Viral Replication. J. Biol. Chem. 2007, 282 (8), 5888–5898. https://doi.org/10.1074/jbc.M611145200.

(16) Vance, L. M.; Moscufo, N.; Chow, M.; Heinz, B. A. Poliovirus 2C Region Functions during Encapsidation of Viral RNA. J. Virol. 1997, 71 (11), 8759–8765. https://doi.org/10.1128/JVI.71.11.8759-8765.1997.

(17) Wang, C.; Ma, H.-C.; Wimmer, E.; Jiang, P.; Paul, A. V. A C-Terminal, Cysteine-Rich Site in Poliovirus 2CATPase Is Required for Morphogenesis. J. Gen. Virol. 2014, 95 (Pt 6), 1255–1265. https://doi.org/10.1099/vir.0.062497-0.

(18) Wang, C.; Jiang, P.; Sand, C.; Paul, A. V.; Wimmer, E. Alanine Scanning of Poliovirus 2CATPase Reveals New Genetic Evidence That Capsid Protein/2CATPase Interactions Are Essential for Morphogenesis. J. Virol. 2012, 86 (18), 9964–9975. https://doi.org/10.1128/JVI.00914-12.

(19) Liu, Y.; Wang, C.; Mueller, S.; Paul, A. V.; Wimmer, E.; Jiang, P. Direct Interaction between Two Viral Proteins, the Nonstructural Protein 2CATPase and the Capsid Protein VP3, Is Required for Enterovirus Morphogenesis. PLoS Pathog. 2010, 6 (8), e1001066. https://doi.org/10.1371/journal.ppat.1001066.

(20) Verlinden, Y.; Cuconati, A.; Wimmer, E.; Rombaut, B. The Antiviral Compound 5-(3,4-Dichlorophenyl) Methylhydantoin Inhibits the Post-Synthetic Cleavages and the Assembly of Poliovirus in a Cell-Free System. Antiviral Res. 2000, 48 (1), 61–69. https://doi.org/10.1016/S0166-3542(00)00119-4.

(21) Suhy, D. A.; Giddings, T. H.; Kirkegaard, K. Remodeling the Endoplasmic Reticulum by Poliovirus Infection and by Individual Viral Proteins: An Autophagy-Like Origin for Virus-Induced Vesicles. J. Virol. 2000, 74 (19), 8953–8965. https://doi.org/10.1128/JVI.74.19.8953-8965.2000.

(22) Teterina, N. L.; Gorbalenya, A. E.; Egger, D.; Bienz, K.; Ehrenfeld, E. Poliovirus 2C Protein Determinants of Membrane Binding and Rearrangements in Mammalian Cells. J. Virol. 1997, 71 (12), 8962–8972. https://doi.org/10.1128/jvi.71.12.8962-8972.1997.

(23) Aldabe, R.; Carrasco, L. Induction of Membrane Proliferation by Poliovirus Proteins 2C and 2BC. Biochem. Biophys. Res. Commun. 1995, 206 (1), 64–76. https://doi.org/10.1006/bbrc.1995.1010.

(24) Cho, M. W.; Teterina, N.; Egger, D.; Bienz, K.; Ehrenfeld, E. Membrane Rearrangement and Vesicle Induction by Recombinant Poliovirus 2C and 2BC in Human Cells. Virology 1994, 202 (1), 129–145. https://doi.org/10.1006/viro.1994.1329.

(25) Li, J. P.; Baltimore, D. An Intragenic Revertant of a Poliovirus 2C Mutant Has an Uncoating Defect. J. Virol. 1990, 64 (3), 1102–1107.

(26) Paul, A. V.; Molla, A.; Wimmer, E. Studies of a Putative Amphipathic Helix in the N-Terminus of Poliovirus Protein 2C. Virology 1994, 199 (1), 188–199. https://doi.org/10.1006/viro.1994.1111.

(27) Wang, S.-H.; Wang, K.; Zhao, K.; Hua, S.-C.; Du, J. The Structure, Function, and Mechanisms of Action of Enterovirus Non-Structural Protein 2C. Front. Microbiol. 2020, 11.

(28) Chen, P.; Li, Z.; Cui, S. Chapter Nine - Picornaviral 2C Proteins: A Unique ATPase Family Critical in Virus Replication. In The Enzymes; Cameron, C. E., Arnold, J. J., Kaguni, L. S., Eds.; Viral Replication Enzymes and their Inhibitors Part A; Academic Press, 2021; Vol. 49, pp 235–264. https://doi.org/10.1016/bs.enz.2021.06.008.

(29) Adams, P.; Kandiah, E.; Effantin, G.; Steven, A. C.; Ehrenfeld, E. Poliovirus 2C Protein Forms Homo-Oligomeric Structures Required for ATPase Activity. J. Biol. Chem. 2009, 284 (33), 22012–22021. https://doi.org/10.1074/jbc.M109.031807.

(30) Xia, H.; Wang, P.; Wang, G.-C.; Yang, J.; Sun, X.; Wu, W.; Qiu, Y.; Shu, T.; Zhao, X.; Yin, L.; Qin, C.-F.; Hu, Y.; Zhou, X. Human Enterovirus Nonstructural Protein 2CATPase Functions as Both an RNA Helicase and ATP-Independent RNA Chaperone. PLOS Pathog. 2015, 11 (7), e1005067. https://doi.org/10.1371/journal.ppat.1005067.

(31) Guan, H.; Tian, J.; Qin, B.; Wojdyla, J. A.; Wang, B.; Zhao, Z.; Wang, M.; Cui, S. Crystal Structure of 2C Helicase from Enterovirus 71. Sci. Adv. 2017, 3 (4), e1602573. https://doi.org/10.1126/sciadv.1602573.

(32) Guan, H.; Tian, J.; Zhang, C.; Qin, B.; Cui, S. Crystal Structure of a Soluble Fragment of Poliovirus 2CATPase. PLoS Pathog. 2018, 14 (9), e1007304. https://doi.org/10.1371/journal.ppat.1007304.

(33) Zhang, C.; Yang, F.; Wojdyla, J. A.; Qin, B.; Zhang, W.; Zheng, M.; Cao, W.; Wang, M.; Gao, X.; Zheng, H.; Cui, S. An Anti-Picornaviral Strategy Based on the Crystal Structure of Foot-and-Mouth Disease Virus 2C Protein. Cell Rep. 2022, 40 (1), 111030. https://doi.org/10.1016/j.celrep.2022.111030.

(34) Chen, P.; Wojdyla, J. A.; Colasanti, O.; Li, Z.; Qin, B.; Wang, M.; Lohmann, V.; Cui, S. Biochemical and Structural Characterization of Hepatitis A Virus 2C Reveals an Unusual Ribonuclease Activity on Single-Stranded RNA. Nucleic Acids Res. 2022, gkac671. https://doi.org/10.1093/nar/gkac671.

(35) Wallach, I.; Dzamba, M.; Heifets, A. AtomNet: A Deep Convolutional Neural Network for Bioactivity Prediction in Structure-Based Drug Discovery; arXiv:1510.02855; arXiv, 2015. https://doi.org/10.48550/arXiv.1510.02855.

(36) Hsieh, C.-H.; Li, L.; Vanhauwaert, R.; Nguyen, K. T.; Davis, M. D.; Bu, G.; Wszolek, Z. K.; Wang, X. Miro1 Marks Parkinson’s Disease Subset and Miro1 Reducer Rescues Neuron Loss in Parkinson’s Models. Cell Metab. 2019, 30 (6), 1131-1140.e7. https://doi.org/10.1016/j.cmet.2019.08.023.

(37) Pfister, T.; Wimmer, E. Characterization of the Nucleoside Triphosphatase Activity of Poliovirus Protein 2C Reveals a Mechanism by Which Guanidine Inhibits Poliovirus Replication *. J. Biol. Chem. 1999, 274 (11), 6992–7001. https://doi.org/10.1074/jbc.274.11.6992.

(38) Ulferts, R.; van der Linden, L.; Thibaut, H. J.; Lanke, K. H. W.; Leyssen, P.; Coutard, B.; De Palma, A. M.; Canard, B.; Neyts, J.; van Kuppeveld, F. J. M. Selective Serotonin Reuptake Inhibitor Fluoxetine Inhibits Replication of Human Enteroviruses B and D by Targeting Viral Protein 2C. Antimicrob. Agents Chemother. 2013, 57 (4), 1952–1956. https://doi.org/10.1128/AAC.02084-12.

(39) Hurdiss, D. L.; El Kazzi, P.; Bauer, L.; Papageorgiou, N.; Ferron, F. P.; Donselaar, T.; van Vliet, A. L. W.; Shamorkina, T. M.; Snijder, J.; Canard, B.; Decroly, E.; Brancale, A.; Zeev-Ben-Mordehai, T.; Förster, F.; van Kuppeveld, F. J. M.; Coutard, B. Fluoxetine Targets an Allosteric Site in the Enterovirus 2C AAA+ ATPase and Stabilizes a Ring-Shaped Hexameric Complex. Sci. Adv. 2022, 8 (1), eabj7615. https://doi.org/10.1126/sciadv.abj7615.

(40) Bauer, L.; Manganaro, R.; Zonsics, B.; Hurdiss, D. L.; Zwaagstra, M.; Donselaar, T.; Welter, N. G. E.; Kleef, R. G. D. M. van; Lopez, M. L.; Bevilacqua, F.; Raman, T.; Ferla, S.; Bassetto, M.; Neyts, J.; Strating, J. R. P. M.; Westerink, R. H. S.; Brancale, A.; Kuppeveld, F. J. M. van. Rational Design of Highly Potent Broad-Spectrum Enterovirus Inhibitors Targeting the Nonstructural Protein 2C. PLOS Biol. 2020, 18 (11), e3000904. https://doi.org/10.1371/journal.pbio.3000904.

(41) Ulferts, R.; de Boer, S. M.; van der Linden, L.; Bauer, L.; Lyoo, H. R.; Maté, M. J.; Lichière, J.; Canard, B.; Lelieveld, D.; Omta, W.; Egan, D.; Coutard, B.; van Kuppeveld, F. J. M. Screening of a Library of FDA-Approved Drugs Identifies Several Enterovirus Replication Inhibitors That Target Viral Protein 2C. Antimicrob. Agents Chemother. 2016, 60 (5), 2627–2638. https://doi.org/10.1128/AAC.02182-15.

(42) Shimizu, H.; Agoh, M.; Agoh, Y.; Yoshida, H.; Yoshii, K.; Yoneyama, T.; Hagiwara, A.; Miyamura, T. Mutations in the 2C Region of Poliovirus Responsible for Altered Sensitivity to Benzimidazole Derivatives. J. Virol. 2000, 74 (9), 4146–4154.

(43) Musharrafieh, R.; Zhang, J.; Tuohy, P.; Kitamura, N.; Bellampalli, S. S.; Hu, Y.; Khanna, R.; Wang, J. Discovery of Quinoline Analogs as Potent Antivirals against Enterovirus D68 (EV-D68). J. Med. Chem. 2019, 62 (8), 4074–4090. https://doi.org/10.1021/acs.jmedchem.9b00115.

(44) Musharrafieh, R.; Kitamura, N.; Hu, Y.; Wang, J. Development of Broad-Spectrum Enterovirus Antivirals Based on Quinoline Scaffold. Bioorganic Chem. 2020, 101, 103981. https://doi.org/10.1016/j.bioorg.2020.103981.

(45) Tang, Q.; Xu, Z.; Jin, M.; Shu, T.; Chen, Y.; Feng, L.; Zhang, Q.; Lan, K.; Wu, S.; Zhou, H.-B. Identification of Dibucaine Derivatives as Novel Potent Enterovirus 2C Helicase Inhibitors: In Vitro, in Vivo, and Combination Therapy Study. Eur. J. Med. Chem. 2020, 202, 112310. https://doi.org/10.1016/j.ejmech.2020.112310.

(46) De Palma, A. M.; Heggermont, W.; Lanke, K.; Coutard, B.; Bergmann, M.; Monforte, A.-M.; Canard, B.; De Clercq, E.; Chimirri, A.; Pürstinger, G.; Rohayem, J.; van Kuppeveld, F.; Neyts, J. The Thiazolobenzimidazole TBZE-029 Inhibits Enterovirus Replication by Targeting a Short Region Immediately Downstream from Motif C in the Nonstructural Protein 2C. J. Virol. 2008, 82 (10), 4720–4730. https://doi.org/10.1128/JVI.01338-07.

(47) Hu, Y.; Kitamura, N.; Musharrafieh, R.; Wang, J. Discovery of Potent and Broad-Spectrum Pyrazolopyridine-Containing Antivirals against Enteroviruses D68, A71, and Coxsackievirus B3 by Targeting the Viral 2C Protein. J. Med. Chem. 2021, 64 (12), 8755–8774. https://doi.org/10.1021/acs.jmedchem.1c00758.

(48) Bauer, L.; Manganaro, R.; Zonsics, B.; Strating, J. R. P. M.; El Kazzi, P.; Lorenzo Lopez, M.; Ulferts, R.; van Hoey, C.; Maté, M. J.; Langer, T.; Coutard, B.; Brancale, A.; van Kuppeveld, F. J. M. Fluoxetine Inhibits Enterovirus Replication by Targeting the Viral 2C Protein in a Stereospecific Manner. ACS Infect. Dis. 2019, 5 (9), 1609–1623. https://doi.org/10.1021/acsinfecdis.9b00179.

(49) Lanke, K. H. W.; van der Schaar, H. M.; Belov, G. A.; Feng, Q.; Duijsings, D.; Jackson, C. L.; Ehrenfeld, E.; van Kuppeveld, F. J. M. GBF1, a Guanine Nucleotide Exchange Factor for Arf, Is Crucial for Coxsackievirus B3 RNA Replication. J. Virol. 2009, 83 (22), 11940–11949. https://doi.org/10.1128/JVI.01244-09.

(50) van Kuppeveld, F. J.; Galama, J. M.; Zoll, J.; Melchers, W. J. Genetic Analysis of a Hydrophobic Domain of Coxsackie B3 Virus Protein 2B: A Moderate Degree of Hydrophobicity Is Required for a Cis-Acting Function in Viral RNA Synthesis. J. Virol. 1995, 69 (12), 7782–7790.

(51) Zhang, L.; Lin, D.; Kusov, Y.; Nian, Y.; Ma, Q.; Wang, J.; von Brunn, A.; Leyssen, P.; Lanko, K.; Neyts, J.; de Wilde, A.; Snijder, E. J.; Liu, H.; Hilgenfeld, R. α-Ketoamides as Broad-Spectrum Inhibitors of Coronavirus and Enterovirus Replication: Structure-Based Design, Synthesis, and Activity Assessment. J. Med. Chem. 2020, 63 (9), 4562–4578. https://doi.org/10.1021/acs.jmedchem.9b01828.

(52) Herod, M. R.; Gold, S.; Lasecka-Dykes, L.; Wright, C.; Ward, J. C.; McLean, T. C.; Forrest, S.; Jackson, T.; Tuthill, T. J.; Rowlands, D. J.; Stonehouse, N. J. Genetic Economy in Picornaviruses: Foot-and-Mouth Disease Virus Replication Exploits Alternative Precursor Cleavage Pathways. PLOS Pathog. 2017, 13 (10), e1006666. https://doi.org/10.1371/journal.ppat.1006666.

(53) Herod, M. R.; Ferrer-Orta, C.; Loundras, E.-A.; Ward, J. C.; Verdaguer, N.; Rowlands, D. J.; Stonehouse, N. J. Both Cis and Trans Activities of Foot-and-Mouth Disease Virus 3D Polymerase Are Essential for Viral RNA Replication. J. Virol. 2016, 90 (15), 6864–6883. https://doi.org/10.1128/JVI.00469-16.

(54) Loundras, E.-A.; Herod, M. R.; Harris, M.; Stonehouse, N. J. Foot-and-Mouth Disease Virus Genome Replication Is Unaffected by Inhibition of Type III Phosphatidylinositol-4-Kinases. J. Gen. Virol. 2016, 97 (9), 2221–2230. https://doi.org/10.1099/jgv.0.000527.

(55) Friesner, R. A.; Murphy, R. B.; Repasky, M. P.; Frye, L. L.; Greenwood, J. R.; Halgren, T. A.; Sanschagrin, P. C.; Mainz, D. T. Extra Precision Glide: Docking and Scoring Incorporating a Model of Hydrophobic Enclosure for Protein-Ligand Complexes. J. Med. Chem. 2006, 49 (21), 6177–6196. https://doi.org/10.1021/jm051256o.

(56) Halgren, T. A.; Murphy, R. B.; Friesner, R. A.; Beard, H. S.; Frye, L. L.; Pollard, W. T.; Banks, J. L. Glide: A New Approach for Rapid, Accurate Docking and Scoring. 2. Enrichment Factors in Database Screening. J. Med. Chem. 2004, 47 (7), 1750–1759. https://doi.org/10.1021/jm030644s.

(57) Friesner, R. A.; Banks, J. L.; Murphy, R. B.; Halgren, T. A.; Klicic, J. J.; Mainz, D. T.; Repasky, M. P.; Knoll, E. H.; Shelley, M.; Perry, J. K.; Shaw, D. E.; Francis, P.; Shenkin, P. S. Glide: A New Approach for Rapid, Accurate Docking and Scoring. 1. Method and Assessment of Docking Accuracy. J. Med. Chem. 2004, 47 (7), 1739–1749. https://doi.org/10.1021/jm0306430.

(58) Farid, R.; Day, T.; Friesner, R. A.; Pearlstein, R. A. New Insights about HERG Blockade Obtained from Protein Modeling, Potential Energy Mapping, and Docking Studies. Bioorg. Med. Chem. 2006, 14 (9), 3160–3173. https://doi.org/10.1016/j.bmc.2005.12.032.

(59) Sherman, W.; Day, T.; Jacobson, M. P.; Friesner, R. A.; Farid, R. Novel Procedure for Modeling Ligand/Receptor Induced Fit Effects. J. Med. Chem. 2006, 49 (2), 534–553. https://doi.org/10.1021/jm050540c.

(60) Sherman, W.; Beard, H. S.; Farid, R. Use of an Induced Fit Receptor Structure in Virtual Screening. Chem. Biol. Drug Des. 2006, 67 (1), 83–84. https://doi.org/10.1111/j.1747-0285.2005.00327.x.

(61) Schrödinger Release 2022-1: Maestro. Available at: https://Www.Schrodinger.Com/Products/Maestro (Accessed: 2022).

(62) Wang, C. Functional Analysis of Poliovirus Protein 2CATPase in Viral RNA Replication and Encapsidation Using Alanine Scanning Mutagenesis. 2011.

(63) Li, Q.; Zheng, Z.; Liu, Y.; Zhang, Z.; Liu, Q.; Meng, J.; Ke, X.; Hu, Q.; Wang, H. 2C Proteins of Enteroviruses Suppress IKKβ Phosphorylation by Recruiting Protein Phosphatase 1. J. Virol. 2016. https://doi.org/10.1128/JVI.03021-15.

(64) Wang, X.; Peng, W.; Ren, J.; Hu, Z.; Xu, J.; Lou, Z.; Li, X.; Yin, W.; Shen, X.; Porta, C.; Walter, T. S.; Evans, G.; Axford, D.; Owen, R.; Rowlands, D. J.; Wang, J.; Stuart, D. I.; Fry, E. E.; Rao, Z. A Sensor-Adaptor Mechanism for Enterovirus Uncoating from Structures of EV71. Nat. Struct. Mol. Biol. 2012, 19 (4), 424–429. https://doi.org/10.1038/nsmb.2255.

(65) Dahmane, S.; Kerviel, A.; Morado, D. R.; Shankar, K.; Ahlman, B.; Lazarou, M.; Altan-Bonnet, N.; Carlson, L.-A. Membrane-Assisted Assembly and Selective Autophagy of Enteroviruses. bioRxiv January 31, 2022, p 2021.10.06.463375. https://doi.org/10.1101/2021.10.06.463375.

(66) Knox, C.; Moffat, K.; Ali, S.; Ryan, M.; Wileman, T. Foot-and-Mouth Disease Virus Replication Sites Form next to the Nucleus and Close to the Golgi Apparatus, but Exclude Marker Proteins Associated with Host Membrane Compartments. J. Gen. Virol. 2005, 86 (Pt 3), 687–696. https://doi.org/10.1099/vir.0.80208-0.

(67) Fujita, K.; Krishnakumar, S. S.; Franco, D.; Paul, A. V.; London, E.; Wimmer, E. Membrane Topography of the Hydrophobic Anchor Sequence of Poliovirus 3A and 3AB Proteins and the Functional Effect of 3A/3AB Membrane Association upon RNA Replication. Biochemistry 2007, 46 (17), 5185–5199. https://doi.org/10.1021/bi6024758.

(68) Xiao, X.; Lei, X.; Zhang, Z.; Ma, Y.; Qi, J.; Wu, C.; Xiao, Y.; Li, L.; He, B.; Wang, J. Enterovirus 3A Facilitates Viral Replication by Promoting Phosphatidylinositol 4-Kinase IIIβ–ACBD3 Interaction. J. Virol. 2017, 91 (19), e00791–17. https://doi.org/10.1128/JVI.00791-17.

(69) Laufman, O.; Perrino, J.; Andino, R. Viral Generated Inter-Organelle Contacts Redirect Lipid Flux for Genome Replication. Cell 2019, 178 (2), 275-289.e16. https://doi.org/10.1016/j.cell.2019.05.030.

(70) van der Schaar, H. M.; Dorobantu, C. M.; Albulescu, L.; Strating, J. R. P. M.; van Kuppeveld, F. J. M. Fat(al) Attraction: Picornaviruses Usurp Lipid Transfer at Membrane Contact Sites to Create Replication Organelles. Trends Microbiol. 2016, 24 (7), 535–546. https://doi.org/10.1016/j.tim.2016.02.017.

(71) Belov, G. A.; van Kuppeveld, F. J. M. Lipid Droplets Grease Enterovirus Replication. Cell Host Microbe 2019, 26 (2), 149–151. https://doi.org/10.1016/j.chom.2019.07.017.

(72) Yin, J.; Liu, Y.; Wimmer, E.; Paul, A. V. Complete Protein Linkage Map between the P2 and P3 Non-Structural Proteins of Poliovirus. J. Gen. Virol. 2007, 88 (Pt 8), 2259–2267. https://doi.org/10.1099/vir.0.82795-0.

(73) Wang, T.; Yu, B.; Lin, L.; Zhai, X.; Han, Y.; Qin, Y.; Guo, Z.; Wu, S.; Zhong, X.; Wang, Y.; Tong, L.; Zhang, F.; Si, X.; Zhao, W.; Zhong, Z. A Functional Nuclear Localization Sequence in the VP1 Capsid Protein of Coxsackievirus B3. Virology 2012, 433 (2), 513–521. https://doi.org/10.1016/j.virol.2012.08.040.

(74) Kingston, N. J.; Grehan, K.; Snowden, J. S.; Shegdar, M.; Fox, H.; Macadam, A. J.; Rowlands, D. J.; Stonehouse, N. J. Development of an Enzyme-Linked Immunosorbent Assay for Detection of the Native Conformation of Enterovirus A71. mSphere 2022, 7 (3), e00088–22. https://doi.org/10.1128/msphere.00088-22.

(75) Adeyemi, O. O.; Joseph C. Ward; Joseph S. Snowden; Natalie J. Kingston; Lee Sherry; Morgan R. Herod; David J. Rowlands; Nicola J. Stonehouse. Functional Advantages of Triplication of the 3B Coding Region of the FMDV Genome. FASEB J. 2021, 35 (2), e21215. https://doi.org/10.1096/fj.202001473RR.

(76) Heringa, J. Local Weighting Schemes for Protein Multiple Sequence Alignment. Comput. Chem. 2002, 26 (5), 459–477. https://doi.org/10.1016/s0097-8485(02)00008-6.

(77) Heringa, J. Two Strategies for Sequence Comparison: Profile-Preprocessed and Secondary Structure-Induced Multiple Alignment. Comput. Chem. 1999, 23 (3–4), 341–364. https://doi.org/10.1016/s0097-8485(99)00012-1.

(78) Simossis, V. A.; Heringa, J. PRALINE: A Multiple Sequence Alignment Toolbox That Integrates Homology-Extended and Secondary Structure Information. Nucleic Acids Res. 2005, 33 (suppl_2), W289–W294. https://doi.org/10.1093/nar/gki390.

(79) Simossis, V. A.; Heringa, J. The PRALINE Online Server: Optimising Progressive Multiple Alignment on the Web. Comput. Biol. Chem. 2003, 27 (4–5), 511–519. https://doi.org/10.1016/j.compbiolchem.2003.09.002.

(80) Sievers, F.; Wilm, A.; Dineen, D.; Gibson, T. J.; Karplus, K.; Li, W.; Lopez, R.; McWilliam, H.; Remmert, M.; Söding, J.; Thompson, J. D.; Higgins, D. G. Fast, Scalable Generation of High-Quality Protein Multiple Sequence Alignments Using Clustal Omega. Mol. Syst. Biol. 2011, 7, 539. https://doi.org/10.1038/msb.2011.75.

