## Supporting information for "Development of Enterovirus anti-viral agents that target the viral 2C protein"

^*^ Author to whom correspondence should be addressed.

**Supplemental Figures**

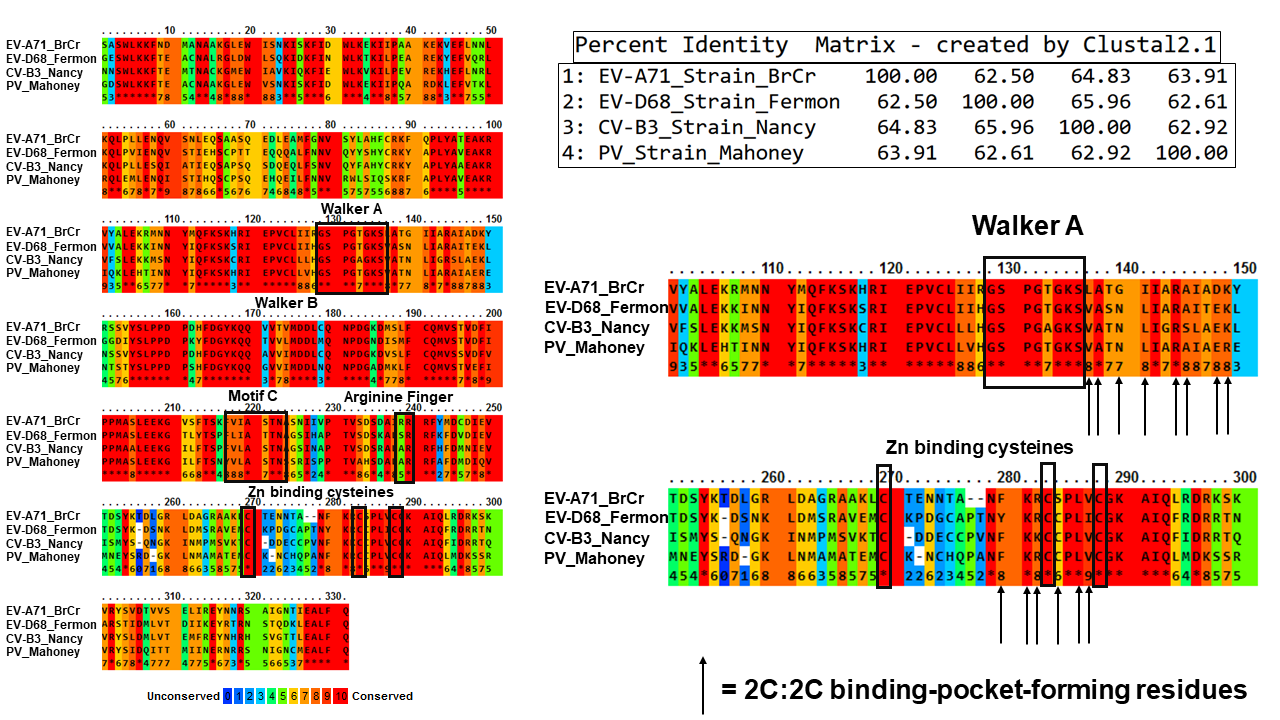

A

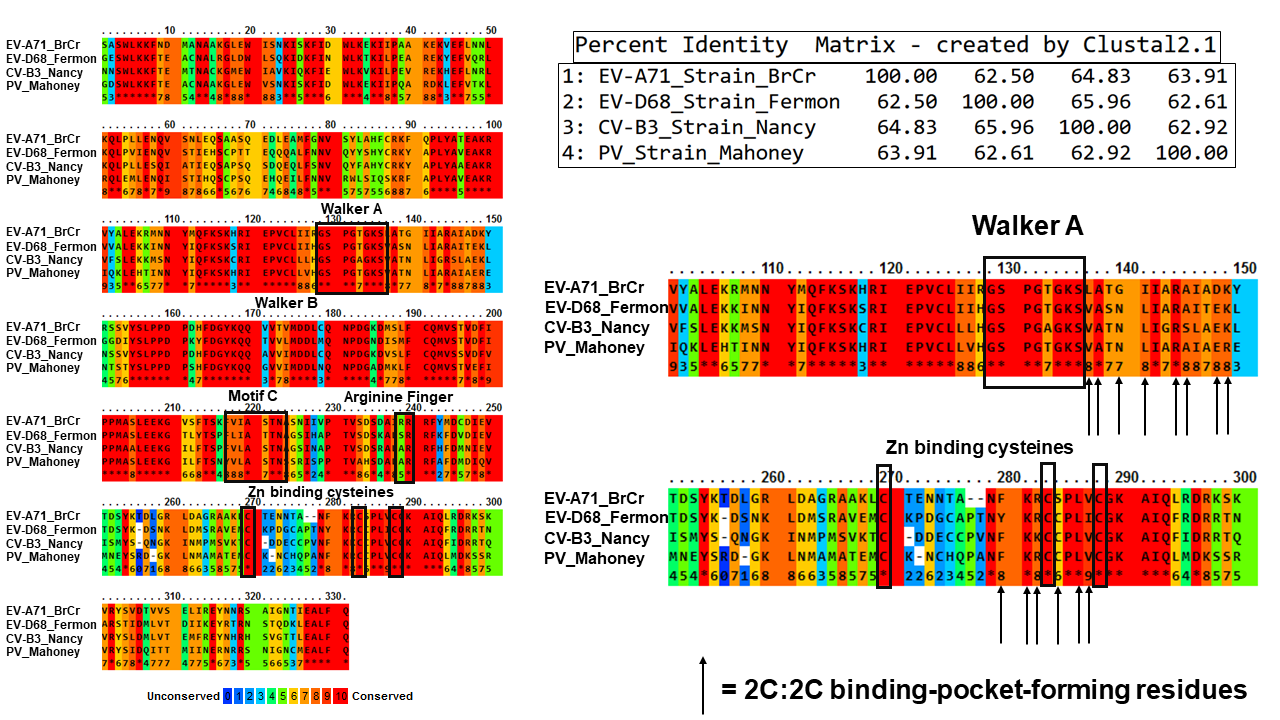

B

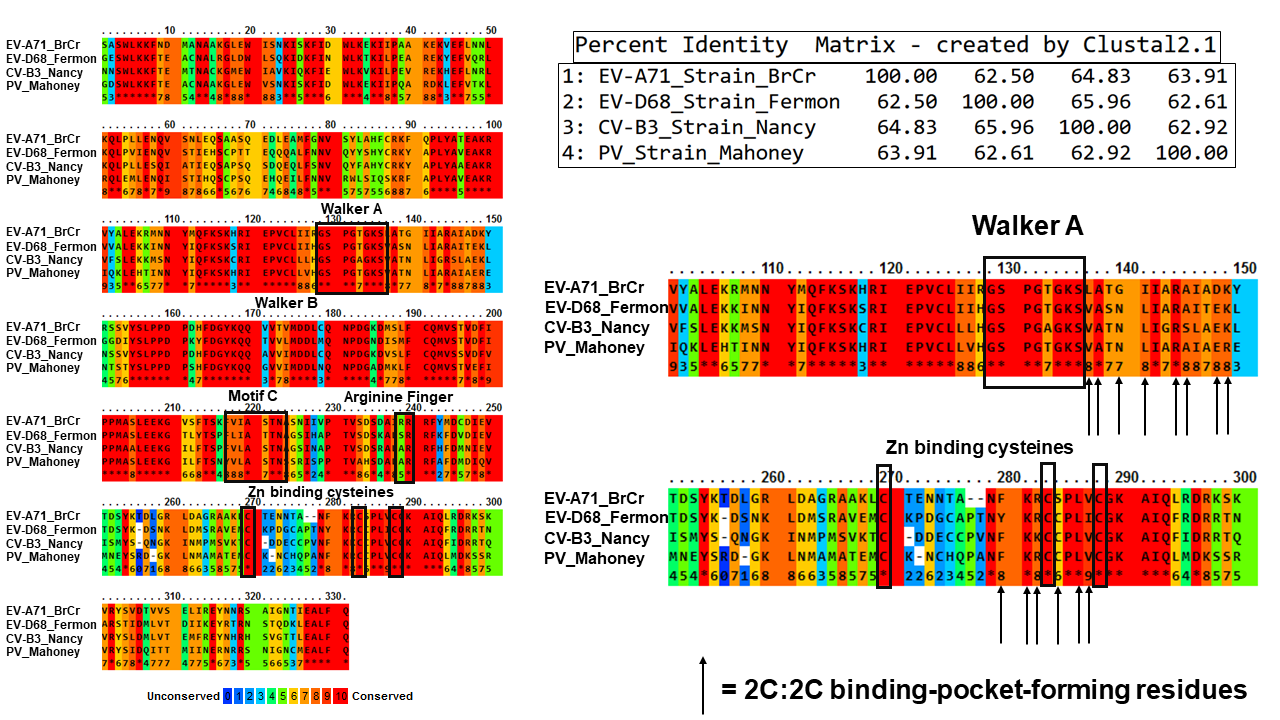

C

**Figure S1. Multiple 2C protein sequence alignment.** A) 2C protein sequences from EV-A71 BrCr strain, EV-D68 Fermon strain, PV Mahoney strain, and Coxsackievirus B3 Nancy strain were aligned using the Praline protein alignment tool and the results for conservation of residues is shown^76–79^. The structural and functional features including Walker A, Walker B, Motif C, Arginine finger, and Zinc binding sites are shown using black boxes. B) The sequences were also aligned using Clustal Omega^80^ and the percent identity matrix was generated and is shown. C) Zoomed in portion of the alignment of A). The 2C:2C binding-pocket-forming residues are highlighted with black arrows.

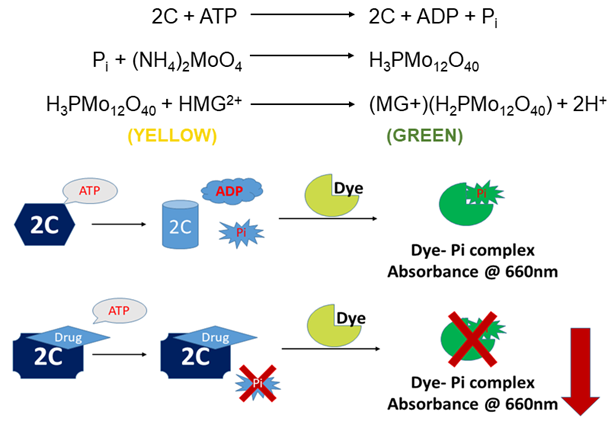

**Figure S2. Schematic of the ATPase assay.** A) 2C is an ATPase and hydrolyzes ATP into ADP and free orthophosphate. This orthophosphate then reacts with the molybdate to form the phosphomolybdate. This complex then reacts with malachite green resulting in a green-colored complex that can be quantified by determining the absorbance at 660 nm. This absorbance reading is a direct correlation to the amount of free phosphate produced as a result of the enzymatic ATPase activity of 2C. Inhibition of 2C by the drug will result in a decreased amount of orthophosphate that will eventually result in a reduced absorbance reading.

**Table S1. Anti-viral activity of the six active compounds against EV-A71.** The assay was performed three times from which the 95 % confidence interval range for IC_50_ was calculated using profile likelihood asymmetrical confidence intervals.

| **Compound Name** | **IC_50_ (µM)** | **95 % confidence interval range (µM)** | **CC_50_ (µM)** | **Selectivity Index** |
| --- | --- | --- | --- | --- |
| SJW-2C-182 | 5 | 2.661 to 7.907 | 306.2 | 61.24 |
| SJW-2C-183 | 16.98 | 15.08 to 19.58 | 85.36 | 5.03 |
| SJW-2C-184 | 2.9 | 2.274 to 3.434 | > 200 | > 68.97 |
| SJW-2C-185 | 12.25 | 10.66 to 13.86 | > 200 | > 16.33 |
| SJW-2C-186 | 102.8 | 76.98 to 152.8 | 61.6 | 0.6 |
| SJW-2C-187 | 44.2 | 41.55 to 47.61 | > 200 | > 4.52 |

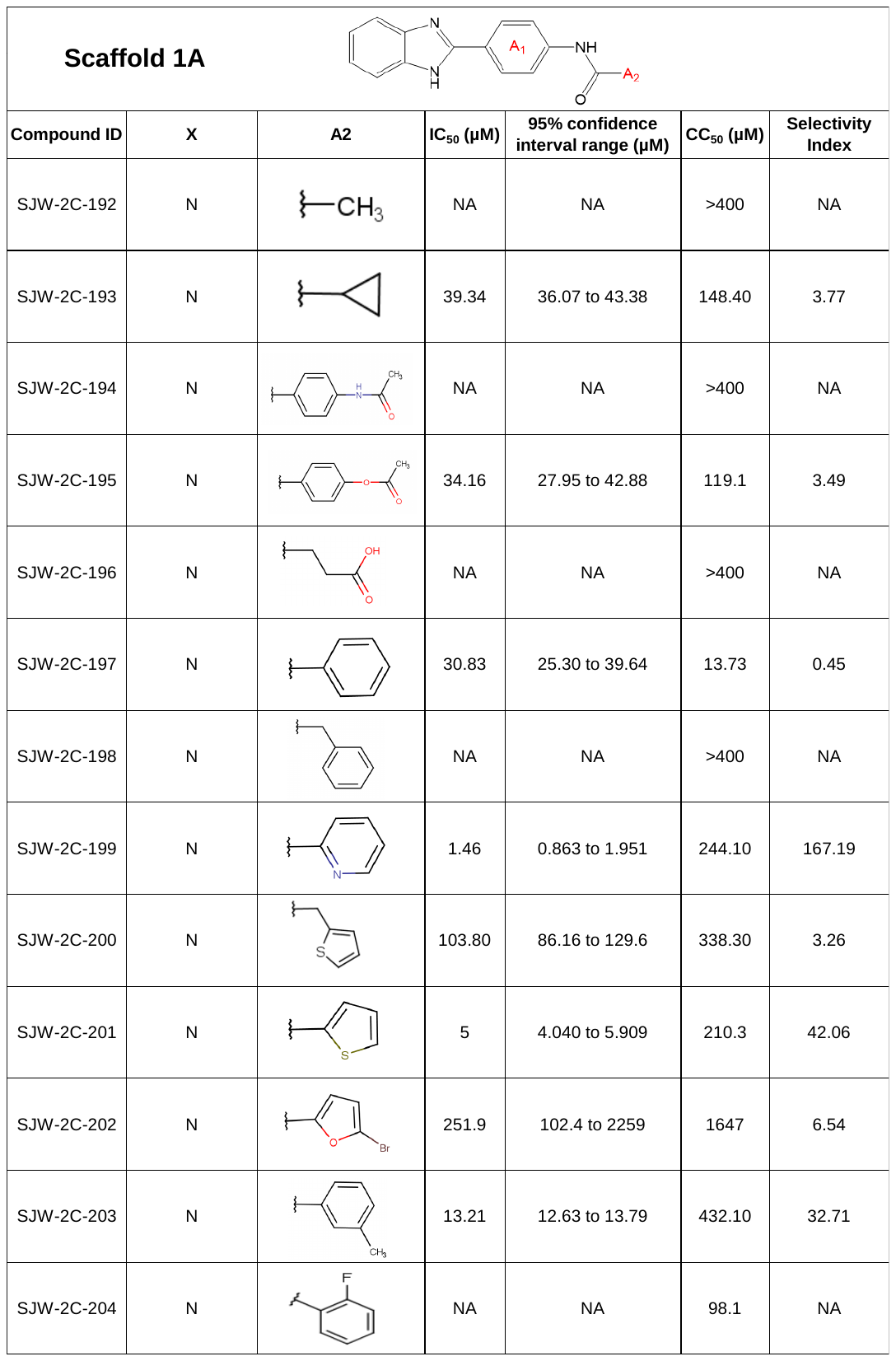
**Table S2. Anti-viral activities of analogs from Scaffold 1A.**

**Table S3. Anti-viral activities of analogs from Scaffold 1B.**

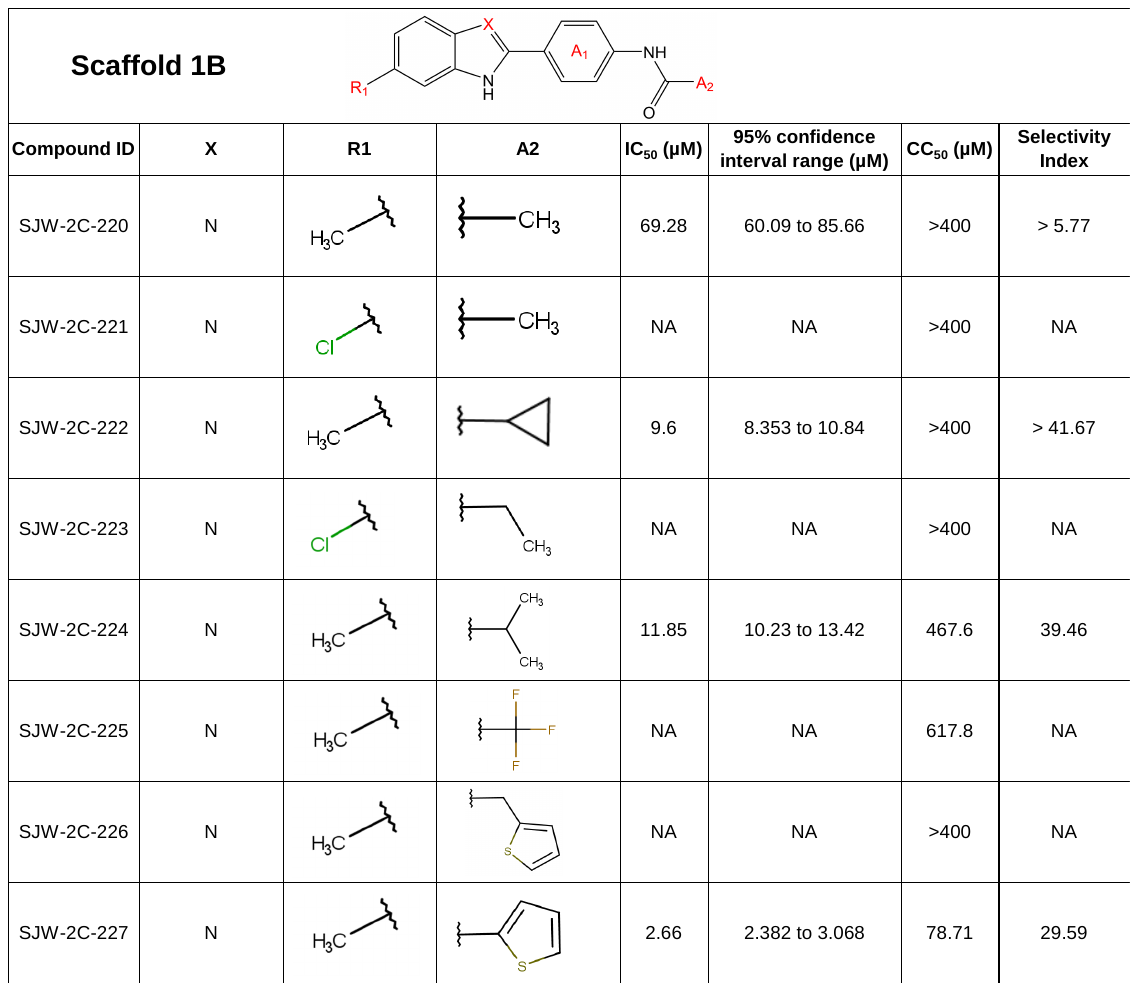

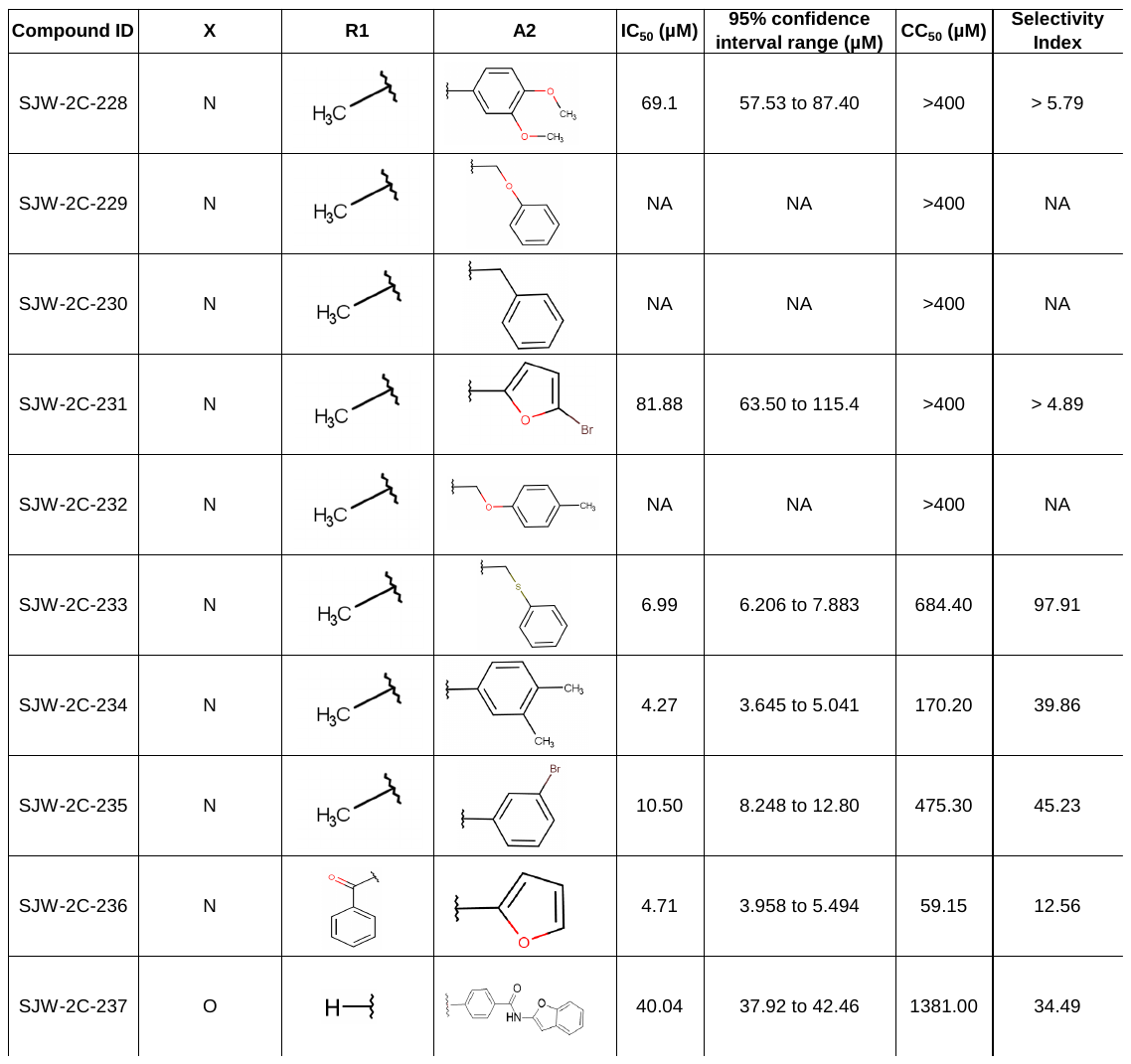

**Table S4. Anti-viral activities of analogs from Scaffold 2.**

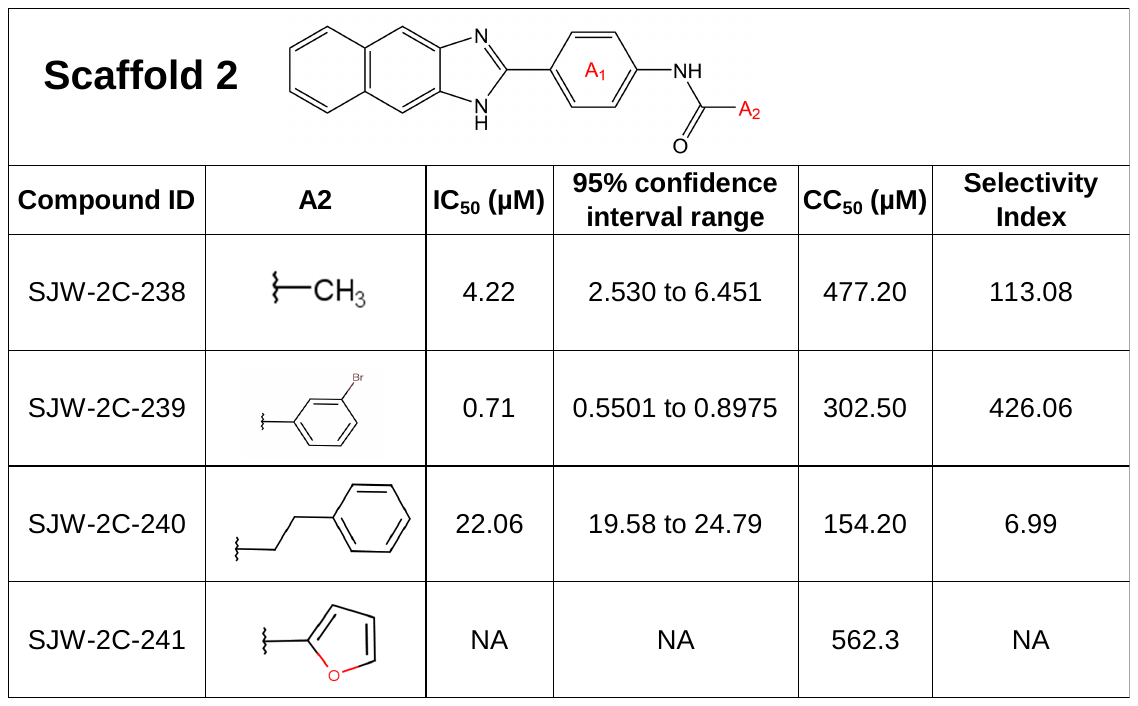

**Table S5. Anti-viral activities of analogs from Scaffold 3.**

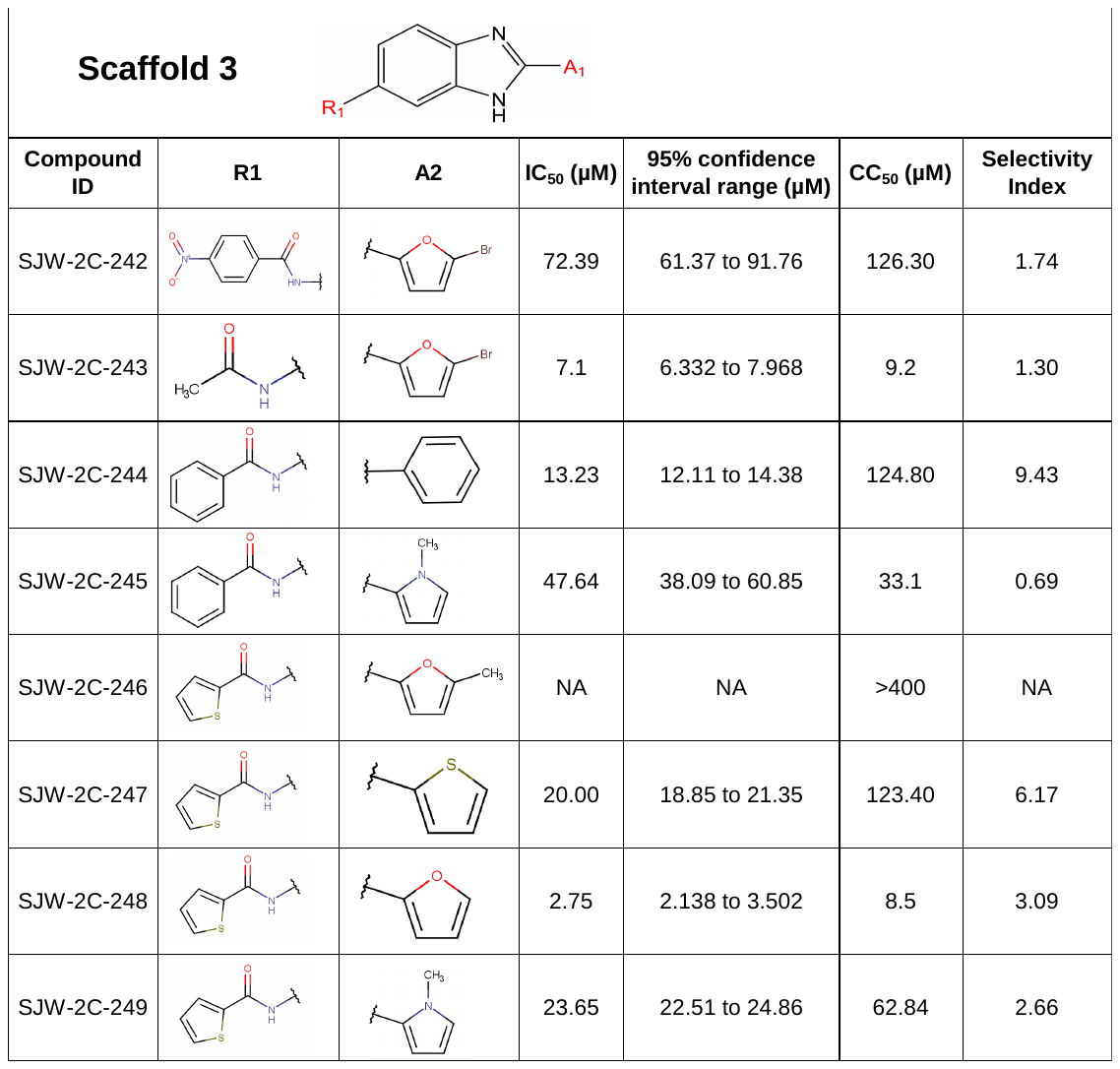

**Table S6. Anti-viral activities of analogs from Scaffold 4.**

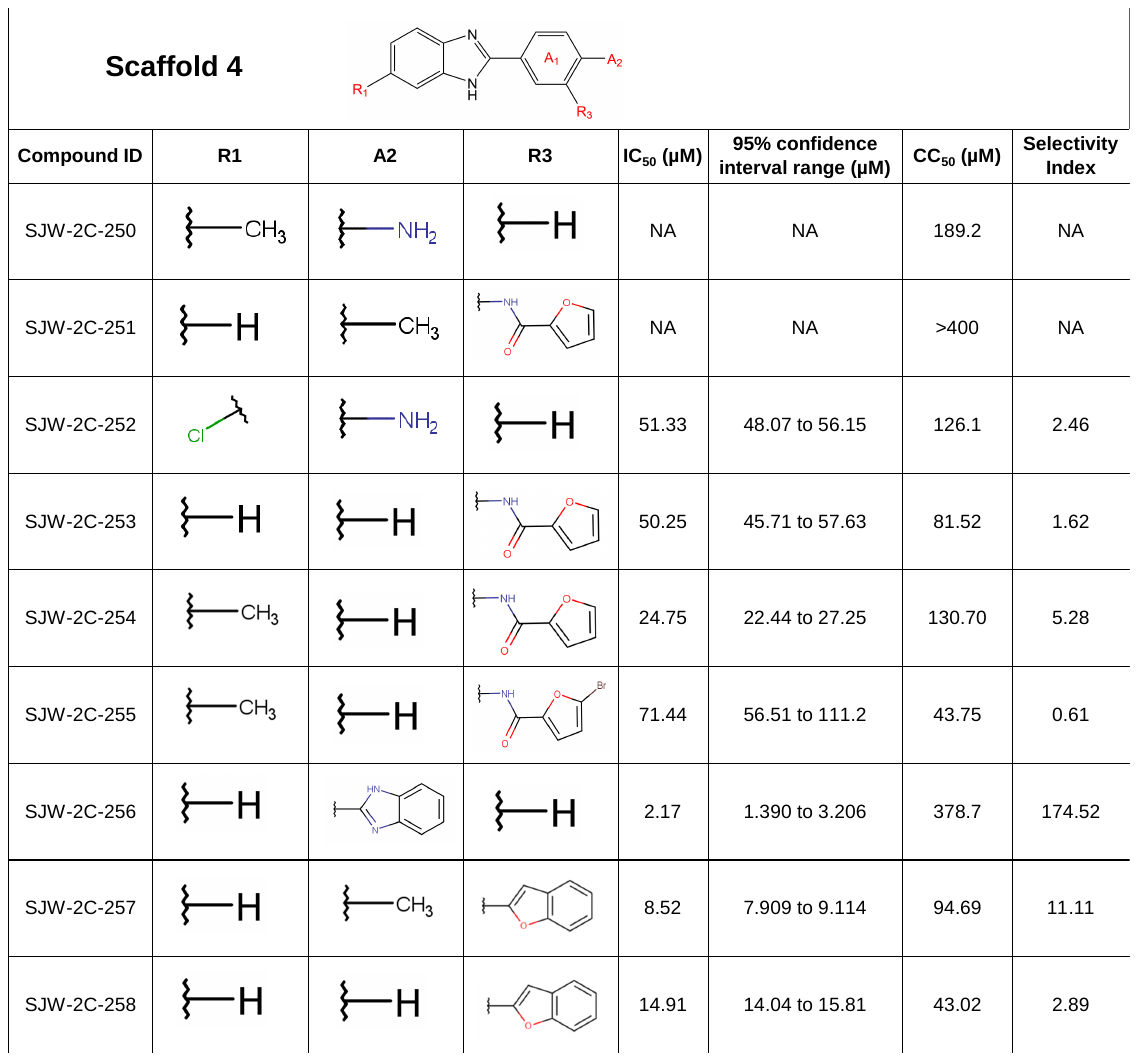

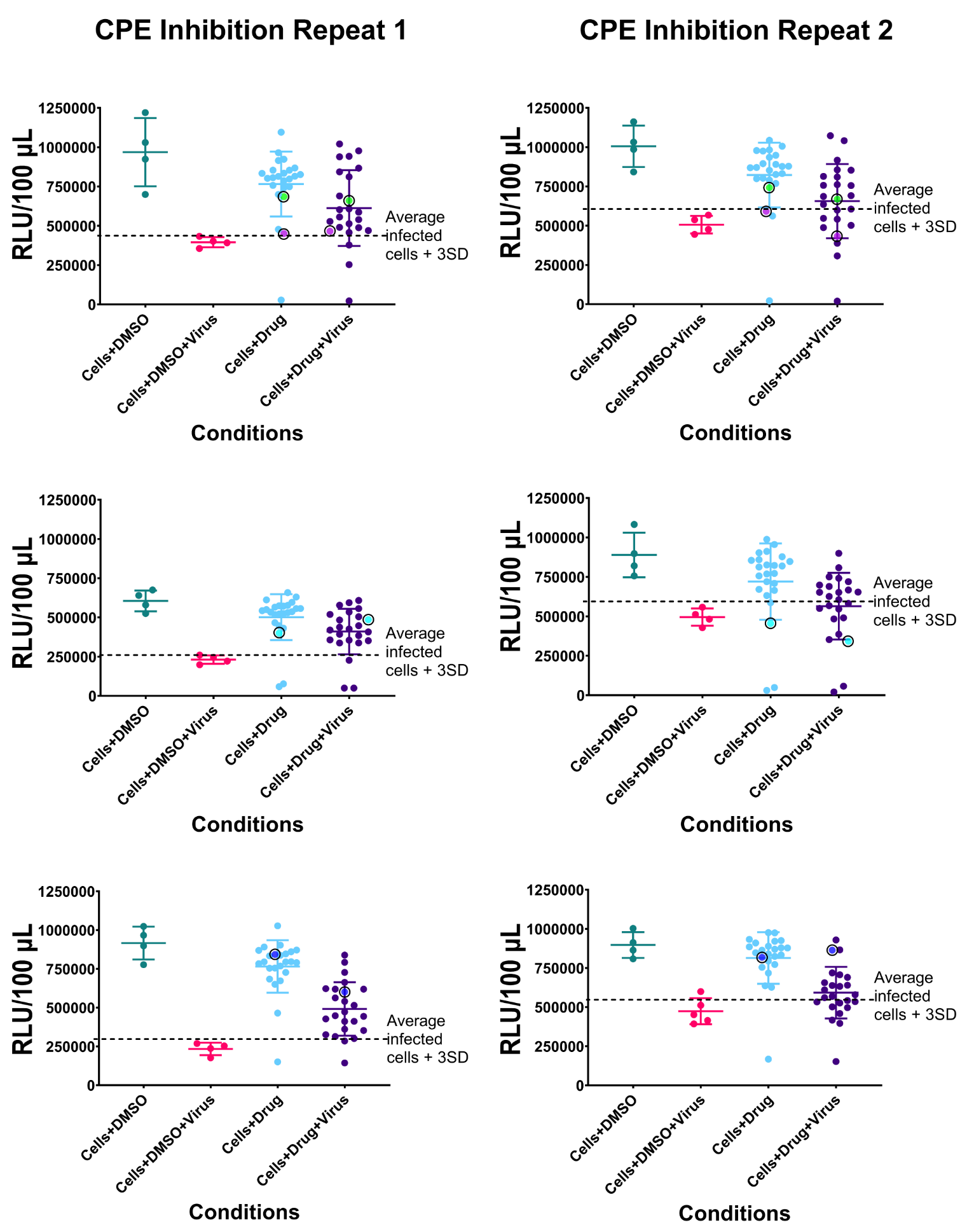

**Figure S3. Compounds tested for inhibition of cytopathic effects (CPE) caused by EV-A71 in Vero cells.** Compounds were screened at a single drug concentration of 50 µM. An assay was performed in duplicate to analyze the effect of drugs on cell viability as well as the cytopathic effect of the virus. The Cell Titer Glo kit from Promega was used to read out the cell viability which directly correlates to the inhibition of CPE caused by EV-A71 and thereby the virus. A threshold of 3x standard deviation (3SD) above the average relative luminescence units (RLU) readout of the infected cells (Cells+DMSO+Virus) was used to identify the anti-viral compounds. SJW-2C-1 is shown as purple circle, SJW-2C-14 is shown as green circle, SJW-2C-33 is shown as cyan circle, and SJW-2C-69 is show as blue circle. The reaction volume was 100 µL.

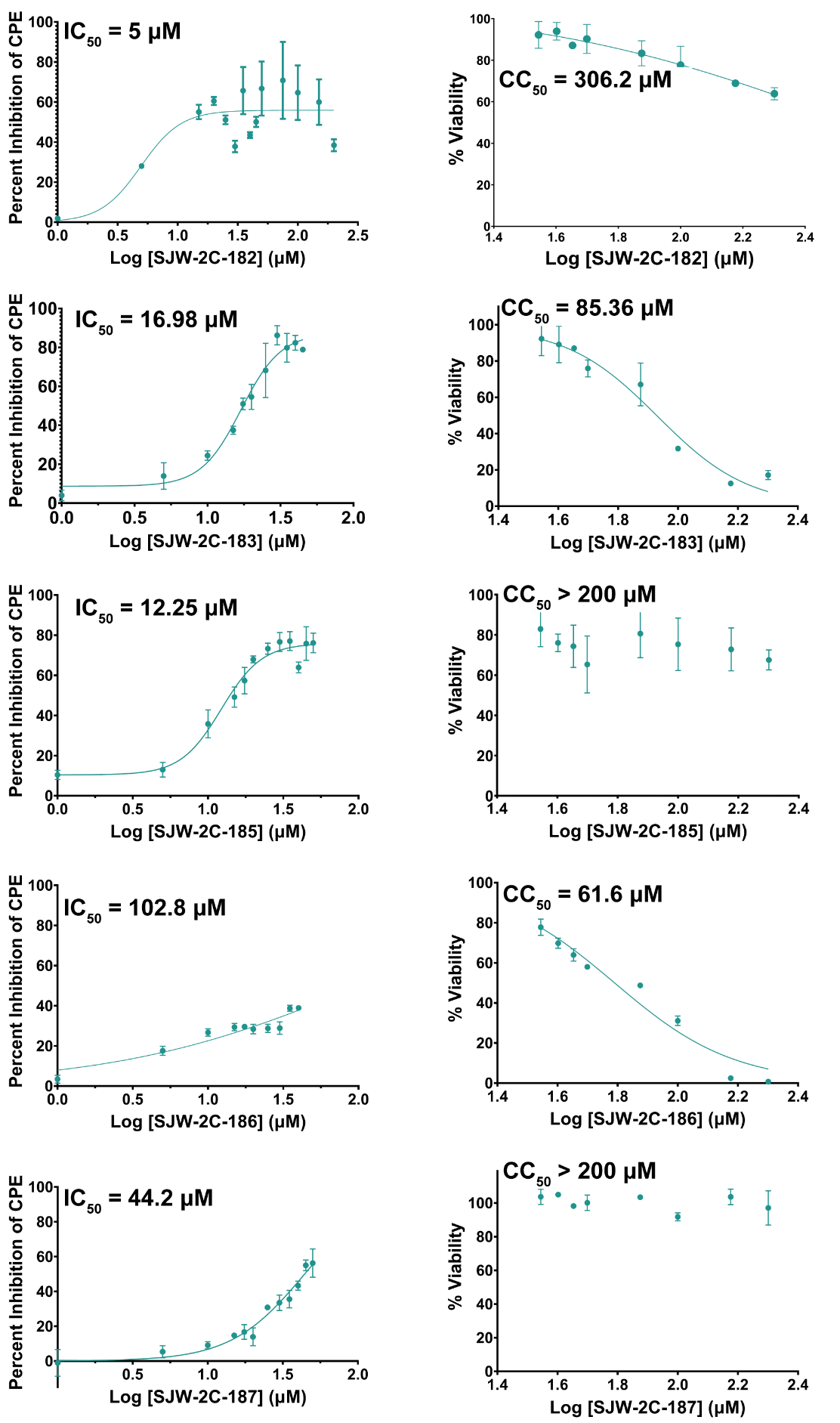

**Figure S4. Dose-response curves for the other five compounds from the structure-activity relationship study of the 45 analogs of SJW-2C-14 and SJW-2C-69.** Dose-response curves for SJW-2C-182, SJW-2C-183, SJW-2C-185, SJW-2C-186 and SJW-2C-187 were also performed to determine their IC_50_ values. The dose range for this assay was 1 µM to 200 µM. Further, a cytotoxicity dose-response curve for each of the compounds were used to calculate their CC_50_ values. The dose range for this assay was 35 µM to 200 µM. Graphed mean and SD, n=3.

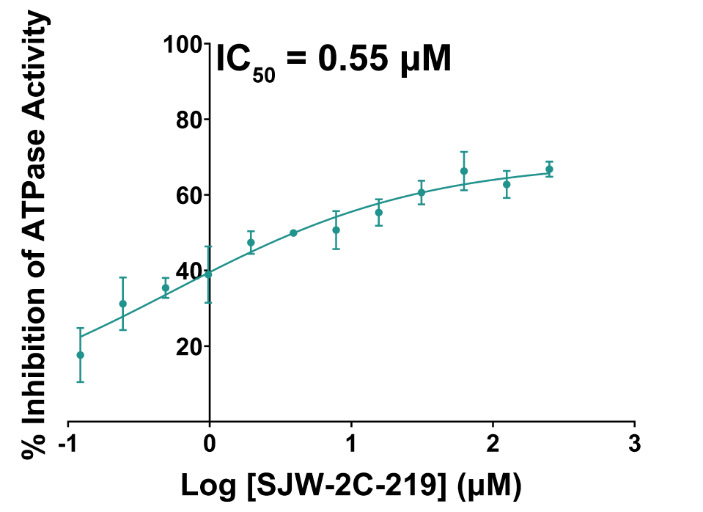

**Figure S5. 2C ATPase inhibition by SJW-2C-219.** Dose-response curve to determine the IC_50_ of Δ2C^116-329^ ATPase inhibition for SJW-2C-219 is shown. The concentration of the compound tested ranged from 0.1 µM to 500 µM. Graphed mean and SD, n=3.

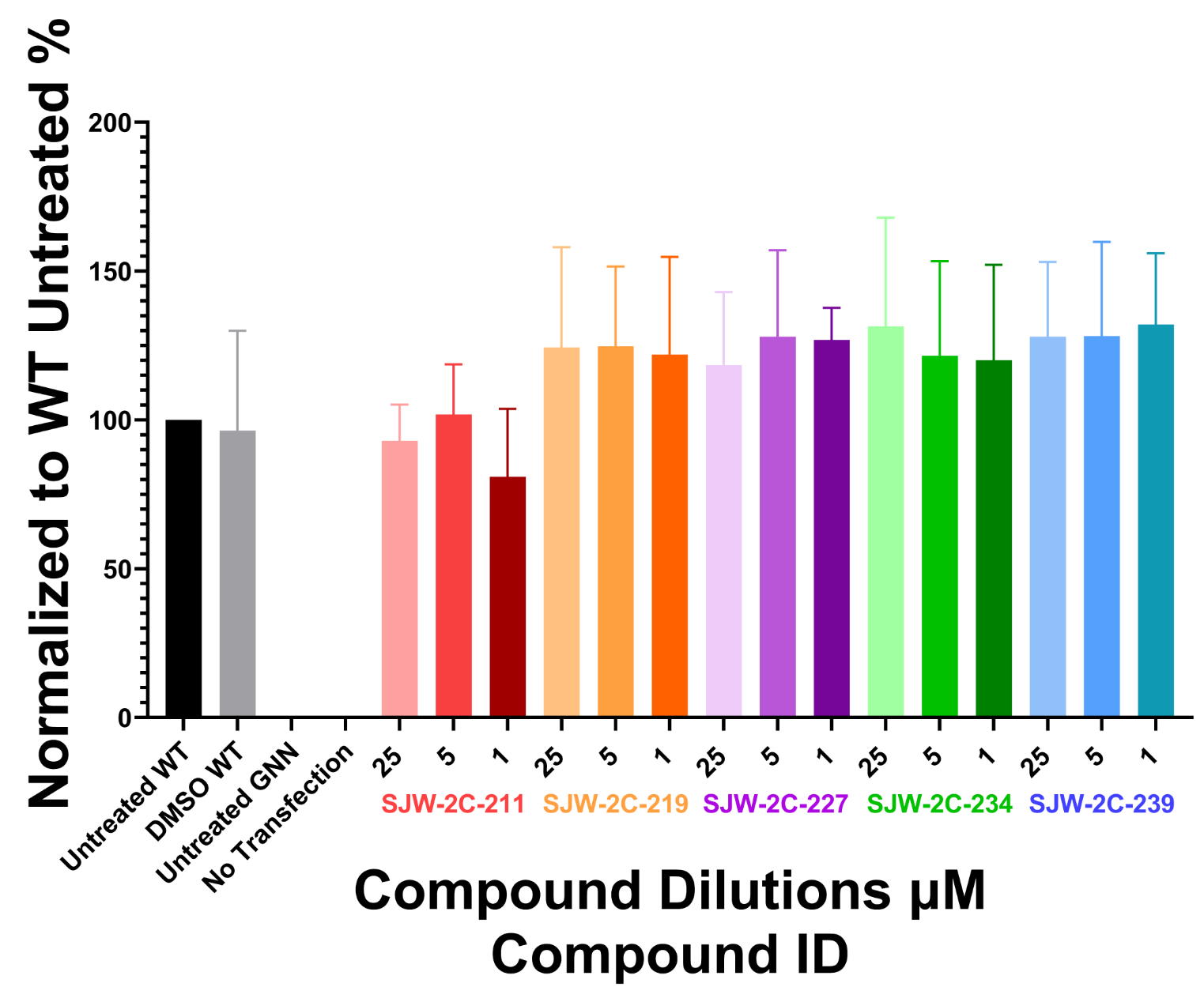

**Figure S6. Effect of SJW-2C-227 on viral RNA replication using the EVA71 replicon system.** Viral RNA levels are quantified upon transfecting Hela cells with the RNA generated from the EV-A71 replicon in the presence of either DMSO (vehicle control) or three different concentrations (25 µM, 5 µM, and 1 µM) of five compounds from this family of compounds, SJW-2C-211, SJW-2C-219, SJW-2C-227, SJW-2C-234 and SJW-2C-239. In this replicon, the P1 capsid encoding region is replaced with the sequence encoding for a fluorescent reporter protein. After 14 hours post-transfection, the fluorescence from the reporter protein was read. The untreated GNN (replication defective) and no transfection controls were used to remove the background fluorescence and the data were normalized to the untreated WT values. Graphed mean and SEM, n=3 in duplicate.

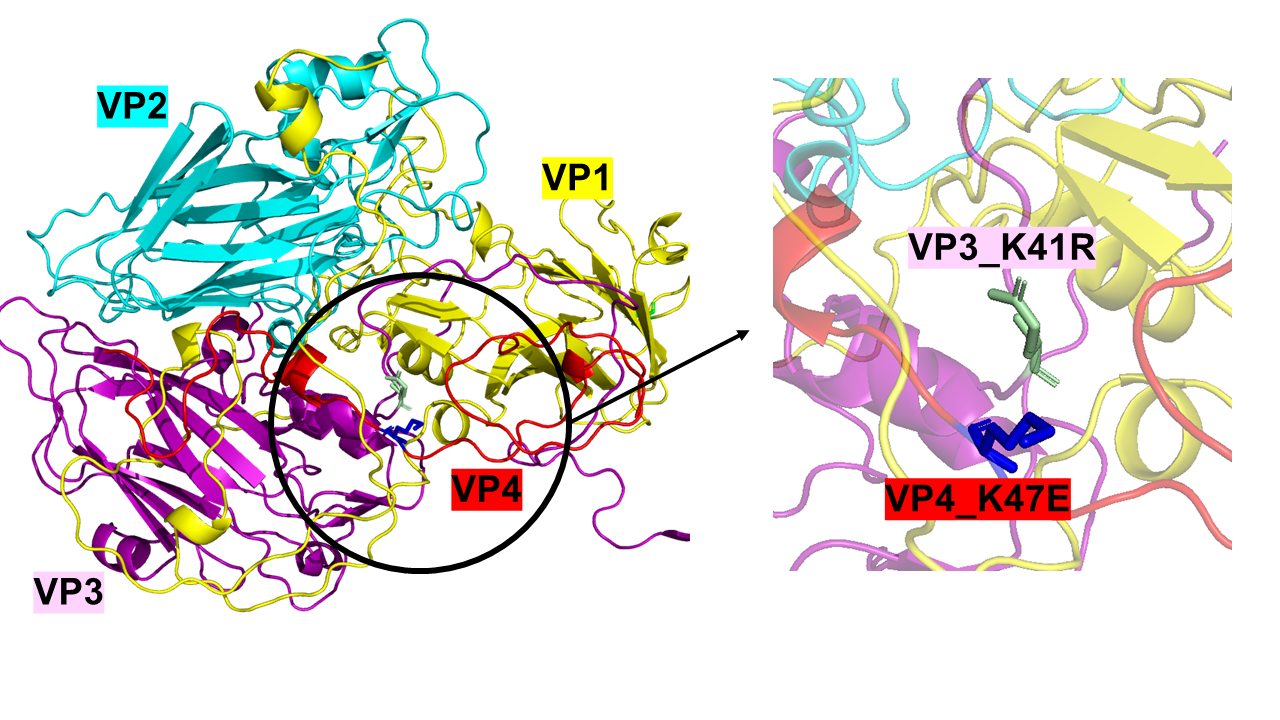

**Figure S7. The proximity of resistant mutant to SJW-2C-227 to encapsidation defect rescue mutant.** Left: EV-A71 protomer showing the four capsid proteins – VP1 in yellow, VP2 in cyan, VP3 in magenta, and VP4 in red (all are rendered as a cartoon) (PDB: 3VBS)^64^. Right: A zoomed-in image showing the proximity of the K47 residue in VP4 to the K41 residue in VP3. K47E is the resistant mutation that was observed in this study to SJW-2C-227 while K41R in PV VP3 is the mutation Wimmer and colleagues observed that rescued an encapsidation defect caused by the mutations K279A/R280A in PV 2C. It is important to note that the residue in EV-A71 VP3 is already an arginine.

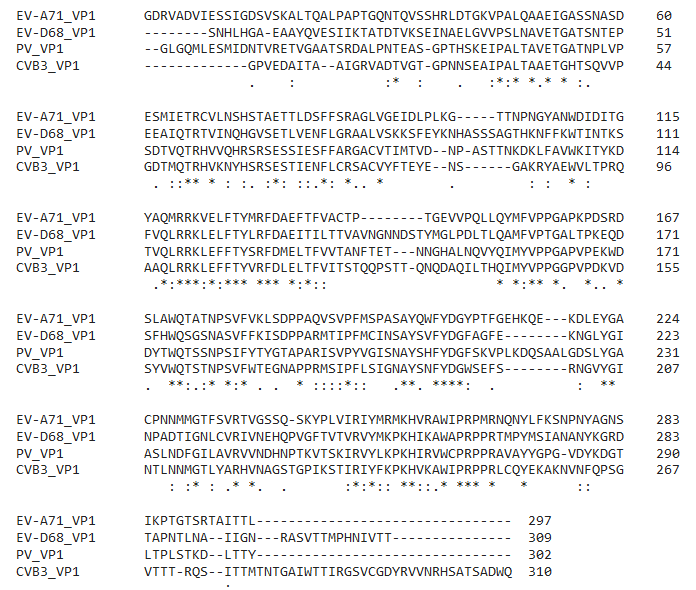

**Figure S8. Alignment of VP1 sequences from EV-A71, EV-D68, PV-1, and CV-B3.** The box highlights the T237 residue in VP1 in EV-A71 and the equivalent residues in the other viruses. Clustal Omega - Multiple Sequence Alignment tool was used for sequence alignment^80^.

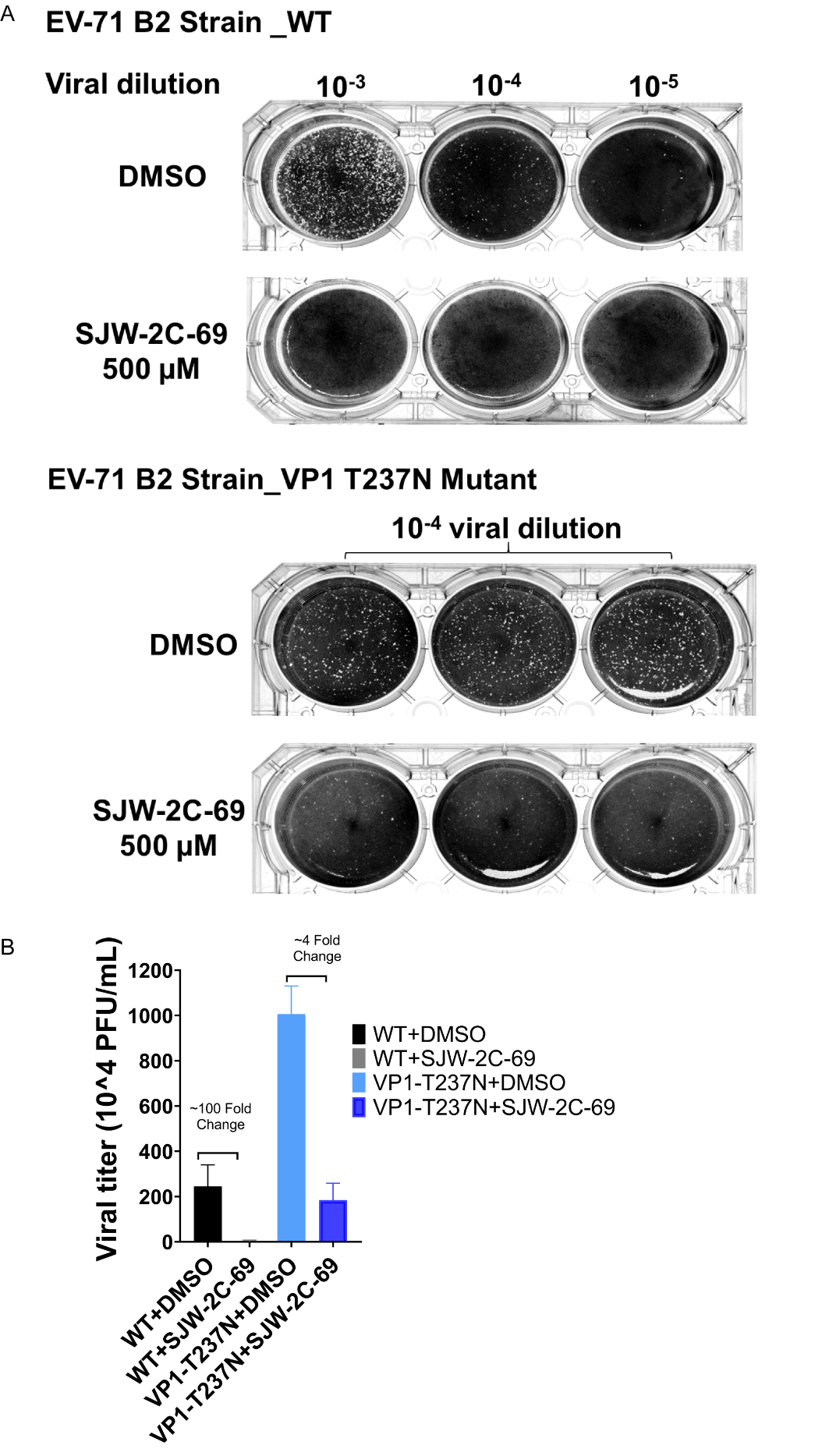

**Figure S9. Validation of the resistant escape mutant.** Plaque assay for viral titer determination for EV-A71 B2 strain wild-type (WT) (top panel) and VP1 T237N (bottom panel) was carried out using Vero cells. Representative plaque assay plates are shown here. Three dilutions of WT virus and one dilution for the mutant virus, when treated with either DMSO (control) or 500 µM of SJW-2C-69 (equal to 10x IC_50_), are shown. After 3 days, plaques were developed, and the viral titer was determined to test the susceptibility of the viruses.
